# Customized and rapid observational delimiting surveys for plant pests based on transect data and scouting

**DOI:** 10.1101/2025.05.08.652837

**Authors:** Barney P. Caton, Godshen R. Pallipparambil, Hui Fang

## Abstract

We have created a complete, standard methodology for designing customized observation-based delimiting surveys for localized incursions of non-endemic plant pests. We propose using transects customized for the species and situation to do the following:

- Collect data for ∼120 detections
- Fit an exponential function to the histogram using an extreme values analysis peaks-over-threshold (EVA POT) technique
- Calculate a percentile-based boundary distance that accounts for two-dimensional (radial) spread
- Use field scouting for verification

We call the approach “Delimitation via Transect Data and Scouting,” or DTDS. In simulations we compared DTDS plans to published survey plans for the Asian longhorned beetle (ALB; *Anoplophora glabripennis* (Motschulsky)), the fungus *Phyllosticta citricarpa* (Guigci), and the tomato brown rugose fruit virus (TBRFV; Tobamovirus). The EVA POT method avoided problems with highly skewed data, and ninety-ninth percentile boundary estimates from it that accounted for radial dispersal always contained the adventive populations. Based on sample sizes, time required, and success (containment) rates the DTDS surveys were efficient and effective. For ALB and TBRFV, DTDS surveys reduced sample sizes and durations by 77 percent or more. For *P. citricarpa*, the DTDS design greatly increased the sample area, but eliminated 6% (scenario 1) and 68% (scenario 2) failure rates in the original plan. The DTDS methodology synthesizes established techniques in a novel way, providing a standardized, clear, and often rapid process for designing customized, transect-based observation-based delimitation surveys that are both efficient and effective. The approach has clear management and financial benefits, and should facilitate quicker, improved decision making.

**Key messages:**

- DTDS is a customizable delimitation survey design method based on transect sampling and scouting
- Transects designed to produce ∼120 positives facilitate efficient sampling of incursion areas
- Boundaries can be accurately determined with uncertainty using extreme values analysis of the distance data
- Case studies demonstrated that using DTDS *always* contained incursions and often greatly reduced survey effort

## Introduction

When new incursions of nonnative plant pests are discovered, determining the spatial extent of populations is usually an important first step in the response (Jang et al. 2014; Kean et al. 2015; van Havre and Whittle 2015). This is done using a delimiting survey, which establishes the boundaries of an area considered to be infested by or free from a pest (IPPC 2016). The goals of delimitation are to confirm the presence of the pest population in a given area and to determine the size of the area occupied (van Havre and Whittle 2015). Some pests cannot be trapped but either can be observed directly (e.g., insects, mollusks, weeds) or produce visible symptoms on hosts (some insects and pathogens), making an observation-based survey the favored approach for delimitation. In some cases, the observation will be collecting samples to test for the presence of a pathogen. Thus, observation-based surveys as used here reflects the sampling procedures of whole plant sampling, visual inspection, and sample collection (IPPC 2021). Recent examples of observation-based surveys for pests in the United States of America include the giant African land snail (GALS; *Lissachatina fulica*, (Férussac)) in Florida (Smith et al. 2010), the spotted lanternfly (SLF; *Lycorma delicatula* (White)) (PDA 2022; PPQ 2022a), and beech leaf disease in the Eastern USA (Forest Service 2022).

Few standard methodologies exist for designing observation-based delimiting surveys for plant pests (Hester et al. 2017), especially if customization by pest type or species is considered. Recent delimitation survey plans by the Plant Protection and Quarantine (PPQ) we reviewed often had methodologies that varied from plan to plan, and some contained little detail. Scientific studies have been done on modeling-informed observation-based survey methods (Hauser and McCarthy 2009; Potts et al. 2013; Lázaro et al. 2020) but these methods have limitations (Becker et al. 2022). In addition, acquiring the needed modeling expertise could take time in situations where quick decision making is preferable (Hester et al. 2017; Ahmed et al. 2022). Specialized modeling has been prioritized more highly in recent years (Lázaro et al. 2020) than more practical, generic methods. Koh et al. (2025) and the European Food Safety Authority (EFSA) (2020a) have described dynamic, adaptive delimitation designs with hypergeometric sampling, which can have significant containment failure rates (see below). With very similar designs, Sun et al. (2025) used individual-based modelling and found that the probability of containment was less than 15 percent overall.

One proposed practical “rapid” observation-based delimitation survey method, by Leung et al. (2010), was termed “Approach, Decline, Delimit” (ADD). ADD uses transects to quickly find the epicenter of the population, approximately locate the boundary (Approach), use sampling to define the decreasing population density (Decline), and quantitatively estimate the boundary while accounting for uncertainty (Delimit). In theoretical simulation studies ADD performed better than naïve sampling schemes but did not work well with insufficient data, or for sparse infestations (Leung et al. 2010). Notably, the authors asked for others to find alternate solutions. Like Hester et al. (2017), we thought the ADD approach—especially the use of transects—was helpful but incomplete. Three remaining questions were how to determine transect lengths, how much area to survey, and how to quantitatively set boundaries that adequately account for uncertainty.

Our objective was to solve these issues and create a more standardized method for designing and implementing observation-based delimiting surveys, which would facilitate more accurate and efficient delimitation of the pest population. The method needed to be rapid to maximize response opportunities (van Havre and Whittle 2015). To minimize costs and simplify planning, we also specifically targeted a “one-shot” approach, as opposed to iterative ones (EFSA et al. 2020a). Here we propose a detailed approach, called “Delimitation via Transect Data and Scouting” (DTDS), based on the following solutions to the methodological needs described above:

1. Define transect length based on relevant biological and situational information
2. Use estimated pest density to set the total survey area needed to give 120 detections
3. Use extreme values analysis to fit functions to the data and derive boundaries

We also advocate using scouting to verify the boundary is accurate.

Below we discuss the rationale and evidence for each component in our DTDS approach and explain the design process and implementation. We applied the design scheme to create comparisons to three published observation-based survey plans or activities. We also used hypothetical spatial data for positives to demonstrate the effectiveness of the transect sampling plans and that EVA POT-based boundaries were more accurate than a more basic curve fitting technique.

## Methods

### DTDS design rationale

Below we describe and justify the proposed components in the DTDS approach for designing customized observation-based delimiting surveys. We refer to “positives” as detected pests or infested hosts. Infestation includes infection (IPPC 2016).

Like Leung et al. (2010) we assumed that only the initial detection(s) was known about the pest population, which located the initial epicenter. They suggested that an uncertain epicenter could be more precisely located using two perpendicular transects. This process is little discussed in the literature. Detections—and especially observation-based detections—are statistically most likely to occur where population density is greatest, near the source (Meats 1998a). That should be especially true for less motile organisms such as weeds (e.g., Scott and Batchelor 2014). We have found little evidence that initial observation-based detections often occur relatively far from the actual epicenter. That seems more likely to happen with trapping, where detection trapping densities are sparse (Meats 1998b), and even then differences were often on the order of only a few hundred meters (Meats 1998a). The DTDS method facilitates evaluating the accuracy of the epicenter through distance data curve fitting, and if uncertainty exists, we suggest using scouting or two quick transects (see below).

#### Component 1. Field scouting

Pre-survey pilot studies are recommended (Anderson et al. 1979; Buckland et al. 2007; Bate et al. 2008) but none of the published survey protocols we have reviewed included any scouting. It requires time and effort (i.e., “boots on the ground”) but provides immediate information, which managers can exploit. For both pre- and post-survey scouting, technologies such as unmanned aerial vehicles (UAVs, or “drones”) may be effective for some pest or symptom types (e.g., Kalischuk et al. 2019; James and Bradshaw 2020).

Pre-survey scouting might be used to avoid simple mistakes or correct protocols before losing time to inaccurate or unproductive efforts. It may only require a few hours, or up to one day, and could be useful for estimating pest (or host) densities, identifying the epicenter, informing personnel about the area and habitat types, and determining how much effort may be needed. Options for the investigation include less and more formal (see below) approaches to setting the pre-survey area(s) or time.

Post-survey scouting aims to verify (or disprove) the boundary determination, by inspecting potential habitats outside that area (e.g., Keith 2000). We recommend inspecting defined areas to give a known level of confidence (see below), applied to different portions of the boundary.

Identifying areas where the target species seems more likely to be present (Hauser and McCarthy 2009; Becker et al. 2022) could enhance the activity, and might be facilitated by knowledge gained during the survey (e.g., host distribution, habitat suitability, or wind patterns; Hull-Sanders et al. 2017).

#### Component 2. Transects

Every recent survey by PPQ that we reviewed used area sampling, usually with ad-hoc cell sizes in fields (e.g., PPQ 2018; PPQ 2020b). Some plans used perimeter-sampling methods, which are well suited to estimating dispersal (see below).

Distance sampling techniques are just as effective and much more efficient than area-based methods (e.g., Anderson et al., 1979; Bauer, 1943; Burnham et al., 1980). In distance sampling, surveyors walk multiple lines through the area of interest, recording where targets are found. The transects are likely to be strips or belts (e.g., Herrick et al., 2005): two-dimensional rectangular areas with a length and width. Most published pest or ecological distance surveys have used parallel transects (e.g., Anderson et al. 1979; Buckland et al. 2015) but the technique is flexible. Distance methods work with motile, immotile, and low motility species (Anderson et al., 1979) and can be used with a broad range of species, including plants (e.g., Buckland et al., 2007), pathogens (e.g., García-Guzmán and Dirzo, 2006), and invertebrates (e.g., Møller and Mousseau, 2009). Formal distance methods usually aim to estimate population densities (Buckland et al. 2015) but recording presence/absence (detection/non-detection; occupancy; frequency) is a substantial simplification (e.g., Forest Service, 2003; Rew et al., 2006) that is usually cheaper and more efficient (MacKenzie et al. 2013). Distance sampling is particularly useful with invasive and rare or sparse species (Elzinga and Salzer 1998; Herrick et al. 2005; Henderson 2010; Bried and Pellet 2012; Thompson 2013; Flesch et al. 2019).

#### Component 3. Transect length

In a circular survey area, the defined transect radius (*R*), ideally extends from the epicenter to just beyond the outermost extent of the pest population. In practice, however, *R* is likely to be overestimated due to uncertainty. Because populations are unlikely to be homogeneous (Philippi 2005; McGarvey et al. 2016), methods suggested for defining *R* by Leung et al. (2010) risk ending the transect too soon, since any non-detection ends the survey.

We propose defining *R* based on estimated time since introduction, species biology, and any other relevant, available information. Time since introduction is important but often uncertain. Generally, longer times since introduction will increase *R*; in one scheme, it is the exact product of the annual dispersal distance and the number of years the pest has been spreading (EFSA et al. 2020b). Relevant information could also include the following.

##### Pest (or vector) dispersal ability

If available, the ideal measure may be annual mean distance traveled (e.g., Skarpaas and Shea 2007; Weldon et al. 2014; Caton et al. 2021), but annual maximum dispersal distance (e.g., Weldon et al. 2014; EFSA et al. 2020b) could work as a conservative approach.

##### Host or habitat ecology

Pests may have one or few obligate hosts, perhaps with limited distributions in the area. Transect studies are frequently adapted to oddly-shaped habitats (Buckland et al. 2015).

##### Field scouting results

Surveyors could find areas away from the epicenter where no more pests are easily found, and then set *R* accordingly.

##### Situational information

Local circumstances might dictate expanding or restricting *R*. For example, Kean et al. (2015) describe a delimiting survey done solely in areas that would conclusively determine if eradication was even practicable.

Defining *R* is a critical task but extreme accuracy is not needed, because later steps in DTDS will help correct any problems by adding transects if necessary, by defining the boundary with uncertainty, and through scouting. Koh et al. (2025) describe radius calculations using an estimated spread rate parameter in various functions (e.g., gamma).

#### Component 4. Sample areas

Sampling inspection activities related to biosecurity often involve hypergeometric distributions (e.g., Vose 2000). However, that distribution applies only to *discrete* populations (Vose 2000; Roberts et al. 2015), so using it to estimate *area-based* sample sizes (e.g., EFSA et al. 2020a) is incorrect. Additionally, detection depends directly on the chosen confidence and infestation levels and failures *will occur* even at expected infestation levels (e.g., Koh et al. 2025). Finally, except in very dense infestations, hypergeometric sample sizes likely yield too few positives for curve fitting.

Sample areas (*A*) are correctly based on the Poisson process, which applies to continuous random variables (Green and Young 1993). Poisson processes describe the probability of observing “events” (i.e., positives) over some interval (time or space) (Vose 2000; Roberts et al. 2015). It is particularly useful for sampling detection of rare species. However, like the hypergeometric, Poisson sampling is a presence/absence test, and so it will also likely produce too few positives. Therefore, we recommend using *m* (density) to find *A* that will confidently give 120 positives. The recommended minimum number of detections in distance sampling is 60 (Buckland et al. 2015), while the United States Forest Service (USFS Pacific Northwest Region 2012) recommended 100 positives. Sixty positives performed well in our tests (below), but since the curve fitting excludes data to focus on extreme values, 120 positives will more confidently result in 60+ distance data points. The formula for estimating the required (critical) sample area (*A*_crit_) is:

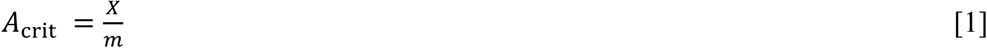

where *X* is the target number of positives. Given *m* = 0.001 per m^2^, *A*_crit_ is 120,000 m^2^ with *X* = 120, whereas the standard Poisson approach would yield a sample area of only 3000 m^2^ (not shown; see Supplementary material on ‘Total transect length calculation’). If *A*_crit_ seems too large to be feasibly surveyed within the program, fewer points might suffice.

Scouting might be used to estimate *m* empirically, using a defined transect length, *L*, and calculating estimated *m* from *L* divided by the number of positives. The choice of *m* could also be arbitrary, based on an infestation level of concern. For example, in the *Phyllosticta citricarpa* case study below they sampled for an infestation rate (*p*_Inf_) of 0.001, or 1 infected tree per 1000 trees (EFSA et al. 2020a). That could be converted to *m* if tree density were known.

#### Component 5. Sampling procedure

This step corresponds to the “Decline” stage of the ADD method (Leung et al. 2010).

The entire transect area, *A*_T_—the product of total transect length (*L*_tot_) and the chosen transect width, *W*_T_—is likely to be inspected, but first we must determine the width and number of transects to be used. We first estimate an initial transect width, *W*_0_, as follows:

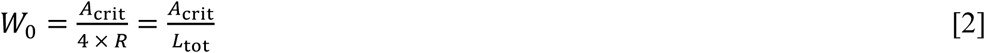

where the 4*R* term assumes that the design has two transects (i.e., *L*_tot_ = 2*R* × 2 transects). Whether *W*_0_ is practicable or not depends on the number of surveyors, and perhaps upon the pest or habitat type (e.g., Henderson 2010). Anderson et al. (1979) argued that *W*_T_ could effectively be unbounded when estimating distances (likely depending on habitat), but widths of 6 m or lower (Herrick et al. 2005; Henderson 2010) seem more practicable here. A single surveyor can likely very effectively cover a 1- to 3-m-wide strip as they proceed; more inspectors can cover wider transects together. In the case studies below, we adopted an upper limit for *W*_T_ for one surveyor of 3 m (0.003 km), and a limit of 6 m (0.006 km) for paired surveyors. For ranges of *A*_crit_ from 0.01 to 0.4 km^2^ and *R* from 0.4 to 6.4 km, about half of the combinations give practicable values of *W*_0_ for one to two surveyors (Fig. 1). Like *R*, *W*_T_ is a factor that could be modified to make a survey more viable, but potential impacts on results should be considered.

**Figure 1.**
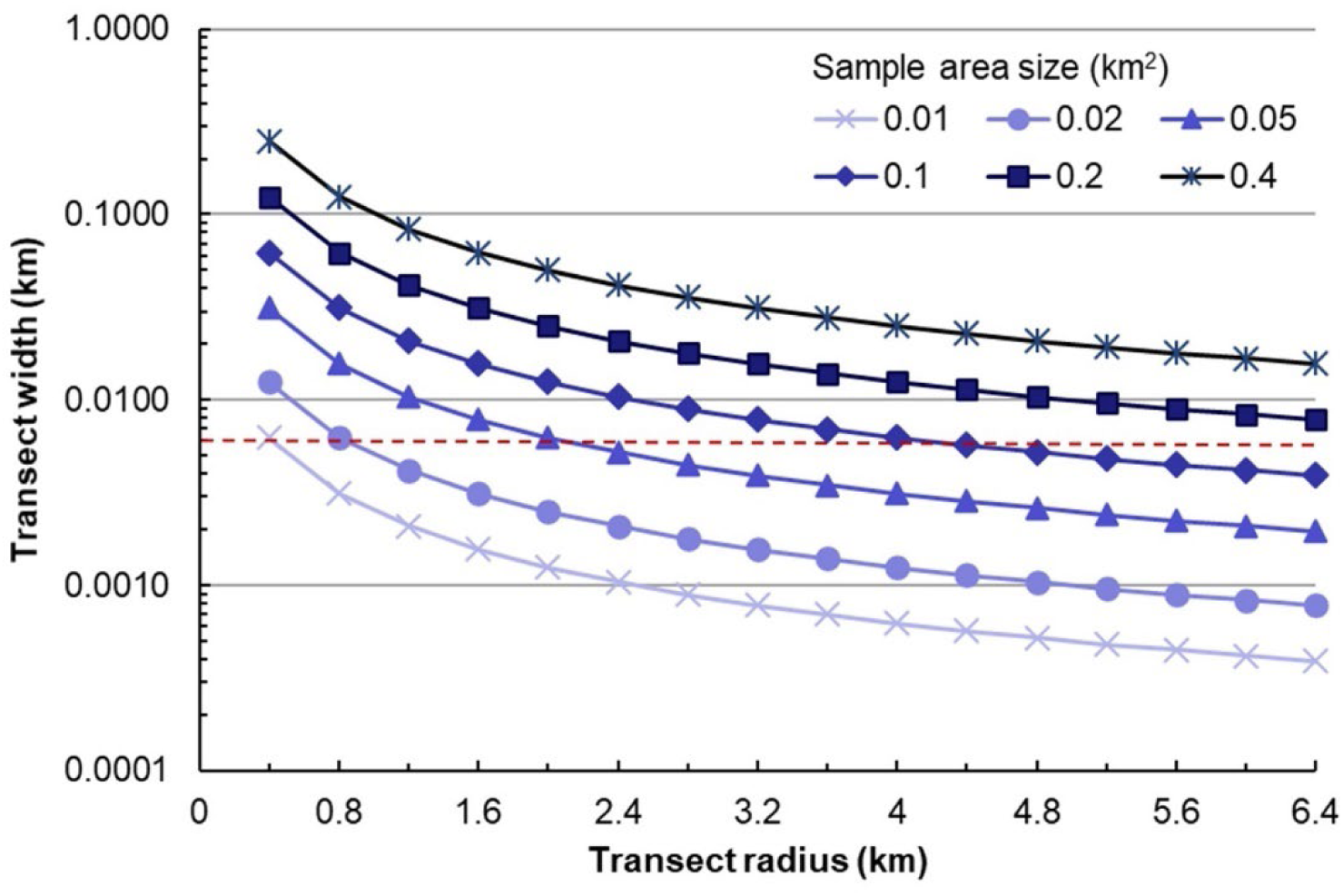
Transect widths as a function of radius. Widths (km) of transect for different recommended sample areas (0.01 to 0.4 km^2^). The dashed red line indicates a threshold value of 6 m for use with two surveyors (see text).

If the value of *W*_0_ indicates that it could be covered by one or two inspectors (i.e., 1 < *W*_0_ < 6), then the proposed specifications should be fine, and the entire area will be sampled. A value of *W*_0_ near zero indicates that *L*_tot_ is relatively very large, and the plan should be modified to randomly sample only part of *A*_T_ (see below).

On the other hand, if *W*_0_ is overly large then more complicated modifications will be needed. Possibilities include adding more transects or lengthening the transect radius. The following function can be used to test adjustments to choose appropriate values for *W*_T_ and the total number of transects used, *N*_T_:

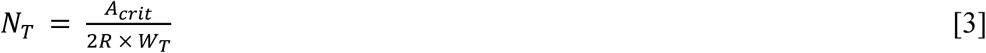

If only a portion of the transect area needs to be inspected, then *W*_T_ should be chosen based on the number of surveyors. Points along the transect (cells) would be randomly chosen for inspection. Using stratified random sampling would help ensure that the full extent of the transects are reasonably covered (Cochran 1977).

#### Component 6. Curve fitting

Frequency distributions of distances traveled or inhabited by organisms from a central point are often termed the “dispersal kernel” (Chapman et al., 2007; Holmes, 1993; Nathan et al., 2012). Dispersal kernels are typically right skewed, because densities are greatest near the epicenter and decrease further away (Weldon et al. 2014).

We fit the dispersal kernel with a two-parameter exponential decay function, which has proven useful and accurate for several species with varying dispersal capabilities (Caton et al. 2021):

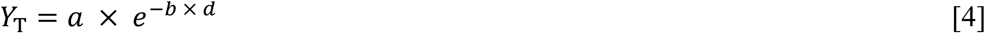

where *Y*_T_ is the dispersal density based on transect (T) data, *a* is the scale parameter (*Y*-intercept), *b* is the (decay) rate, and *d* is the distance. Larger values of *a* and *b* indicate greater densities (probabilities) near the source, and lower densities further from the source.

Detection distance data are likely to vary in their skewness, and dispersal kernel functions are often evaluated and described based on the “fatness” of their distribution tails (Nathan et al. 2012; Bullock et al. 2017). If most detections occur near the epicenter, the fitted boundary may often be underestimated, even with very high percentiles. For example, Fig. 2 shows hypothetical, realistic transect data with most detections near the epicenter but distances up to 1000 m. The 99.9^th^ percentile for the function (equation 4) fitted to the data was only 447 m (not shown).

**Figure 2.**
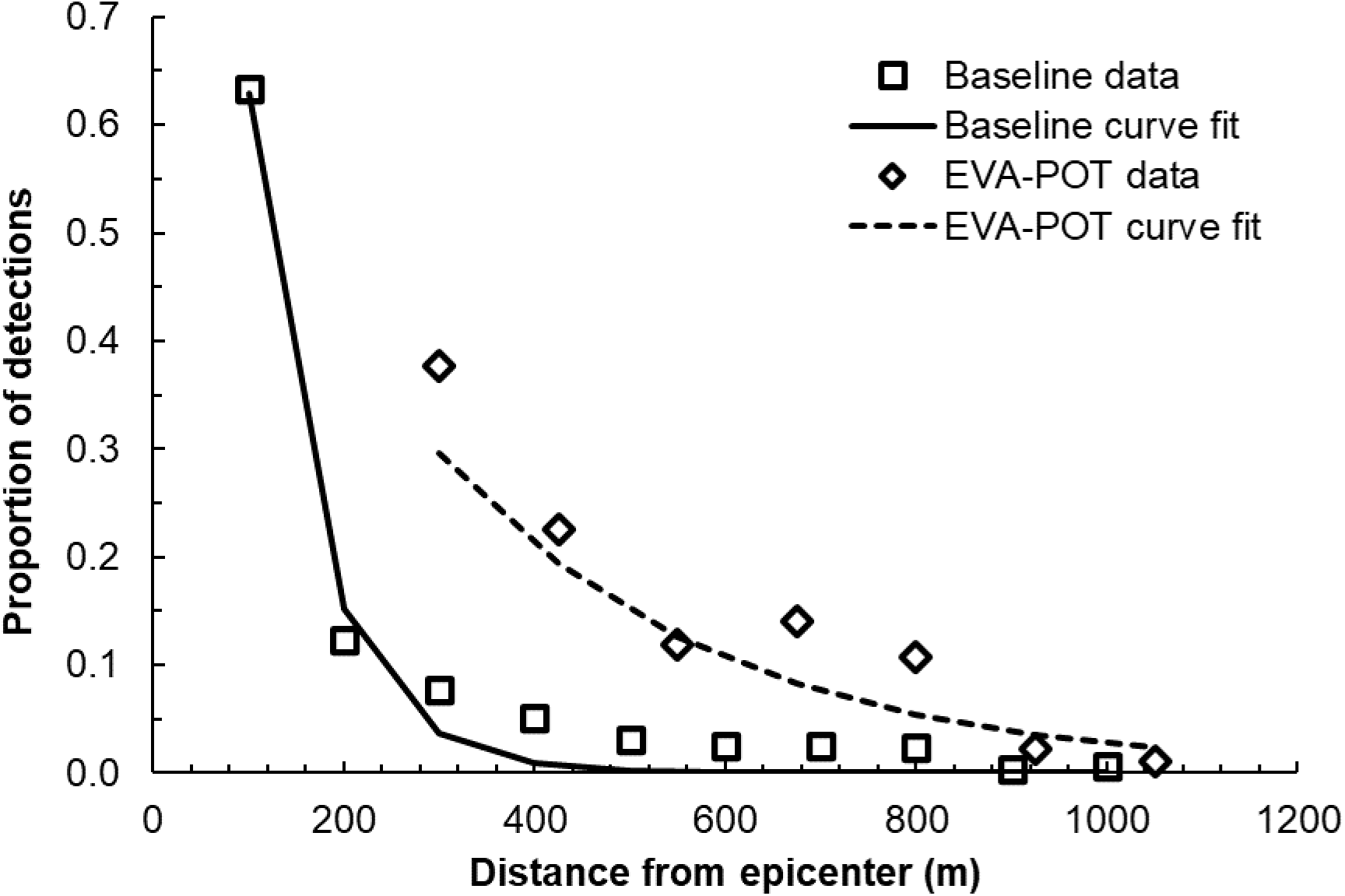
Hypothetical detection data for a non-native invading species that has a true extent of about 1000 m. Shown are observed detections (symbols; plotted on midpoints of bins) and exponential decay curve fits (lines) for either the full dataset (Baseline) or an extreme values analysis peaks-over-threshold (EVA-POT) approach (threshold = 175).

For curve-fitting we used the extreme values analysis peaks-over-threshold technique (EVA POT). This may be an improved method here because it focuses on dataset extremes (Reiss and Thomas 2007; Davison and Huser 2015). EVA has been used extensively to model likelihoods of extreme outcomes, such as wind, rainfall, or flooding events. It has also been applied to ecological processes (e.g., Katz et al. 2005; Jiménez et al. 2011; Carpenter et al. 2018), and we found one application to dispersal (García and Borda-de-Água 2017). The peaks-over-threshold (POT) method seems particularly appropriate for boundary setting and avoiding distributional issues, because only data beyond a certain threshold is included. Applying the EVA POT technique (threshold = 250 m) to the data from above (Fig. 2) yielded a curve fit that gave a 99^th^ percentile boundary of 1320 m, well past the 1000 m extent.

#### Component 7. Boundary estimation

In the ADD method curve fitting and boundary estimation were both contained in the “Delimit” step (Leung et al. 2010). Using a fitted function, the boundary was placed at a distance which gave a statistical expectation that fewer than one cell beyond that distance would be invaded. They seem to be the only researchers who have directly addressed boundary setting.

Regardless of the function chosen, the fitted curve is only a one-dimensional representation of the data. For boundary estimation dispersal in two dimensions needs to be considered (Jones et al. 2009; Nathan et al. 2012), which requires evaluating the radial form of the normalized dispersal kernel (Jones et al. 2009). For the exponential function, the appropriate radial is the Gamma function (*Y*_R_), as follows:

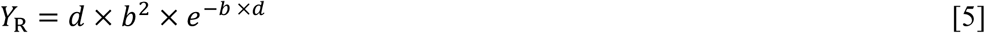

where *b* was fit in equation (4) [note that *a* disappears]. The boundary estimate from equation (5) will be longer and safer but finding its percentiles is not straightforward. Besides using statistical software, which we will not discuss here, an online calculator can be used, such as one by Bognar (2022) that uses α = 2 (constant), fitted *b* (λ in calculator with “rate” option), and the desired *P* (*X* < *x*; e.g., 0.01 for the 99^th^ percentile). Alternately, the percentile (*P*) can be found by trial and error from the following function:

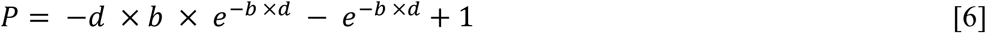

where *d* is the test distance and a result of *P* = 0.990 indicates success for a 99^th^ percentile. Operationally, setting the boundary to the nearest integer should be sufficient.

### Design and Implementation Steps

The proposed DTDS process has six steps, as follows:

1. Scout the area and set the transect radius, *R*.
2. Define the pest infestation density, *m*.
3. Calculate the sample area, *A*_crit_ (equation 1), needed to give 120 positives.
4. Calculate and evaluate the initial transect width, *W*_0_, based on *A*_crit_ and *R* (equation 3).
5. Use *W*_0_ if appropriate (i.e., not too wide), or, if not, modify the number of transects (*N*_T_; equation 4) to get to appropriate values for *W*_T_ and *A*_T._
6. Finalize the design and prepare for field implementation.

The resulting design is then implemented as follows:

1. Lay out and inspect the defined transect areas (and hosts, if relevant) for the target pest (or infested host).
2. Record the distance from the epicenter of each detection (*s*).
3. For insufficient *s* (< 120), lengthen or add transects [repeat Steps 1 and 2]. Move to step 4 when *s* ≈ 120.
4. Convert the data to proportional frequencies as a function of distance.
5. Use standard EVA POT routines to define the threshold, exclude the affected data, and recalculate the histogram.
6. Fit the two-parameter exponential curve (equation 4) to the histogram.
7. Use the rate parameter (*b*) to estimate the 99^th^ percentile (or other desired value) of the radial function (equation 5).
8. Scout areas at and beyond the boundary distance, focusing on suitable habitat and/or hosts.

a. After a thorough search with no pests detected beyond the boundary, set the final boundary.
b. If pests are detected beyond the boundary, go to Step 9.
9. Determine with scouting (or other means) whether the pest population is still localized or more widespread.

a. If the population still seems localized, restart the survey design process at Step 1 with a new (longer) *R*.
b. If the pest population is more wide-spread, stop this process and consider area-wide delimitation methods.

Although not listed above, in-field training and testing of the survey protocols, or a full pilot study (e.g., “test” transects), are recommended before beginning the survey in earnest (Buckland et al. 2015). Tips for maximizing the detectability of target pests can be found in Henderson (2010) and Turgeon et al. (2010).

### Case study specifications

We identified published observation-based delimitation survey plans for three species, all of which had host plants: *Anoplophora glabripennis* (Motschulsky) (Asian longhorned beetle; or ALB), the fungus *Phyllosticta citricarpa* (Guigci), and the tomato brown rugose fruit virus (TBRFV; Tobamovirus). We created customized DTDS plans for each species and used stochastic simulations to compare effort and outcomes for the published and DTDS plans.

Models emulated the survey conditions (e.g., areas, host densities, infestation rates) and survey plan specifications (areas and hosts inspected). Situational details were constant within each case study; only survey specifications differed. The main outputs were the mean number of infected/infested hosts detected, by plan or scenario, the total time required, and the success rate. For the DTDS plans, we used separate stochastic models to produce spatial data from transect sampling, and used that data to evaluate the curve fitting and boundary setting processes. Note that the DTDS plans were originally designed to produce only 60 positives.

#### Asian longhorned beetle (ALB)

##### Published plan description

The delimitation (i.e., “Level 2”) survey plan dictates inspecting all hosts within a radius of 2.4 km (1.5 mi; Table 1), assuming smaller population levels (PPQ 2014). The total area surveyed in one location would therefore be 18.1 km^2^. We used a mean density of 25,000 trees per km^2^ (Dodds and Orwig 2011) for known host species (PPQ 2014) (Table S1).

**Table 1.**
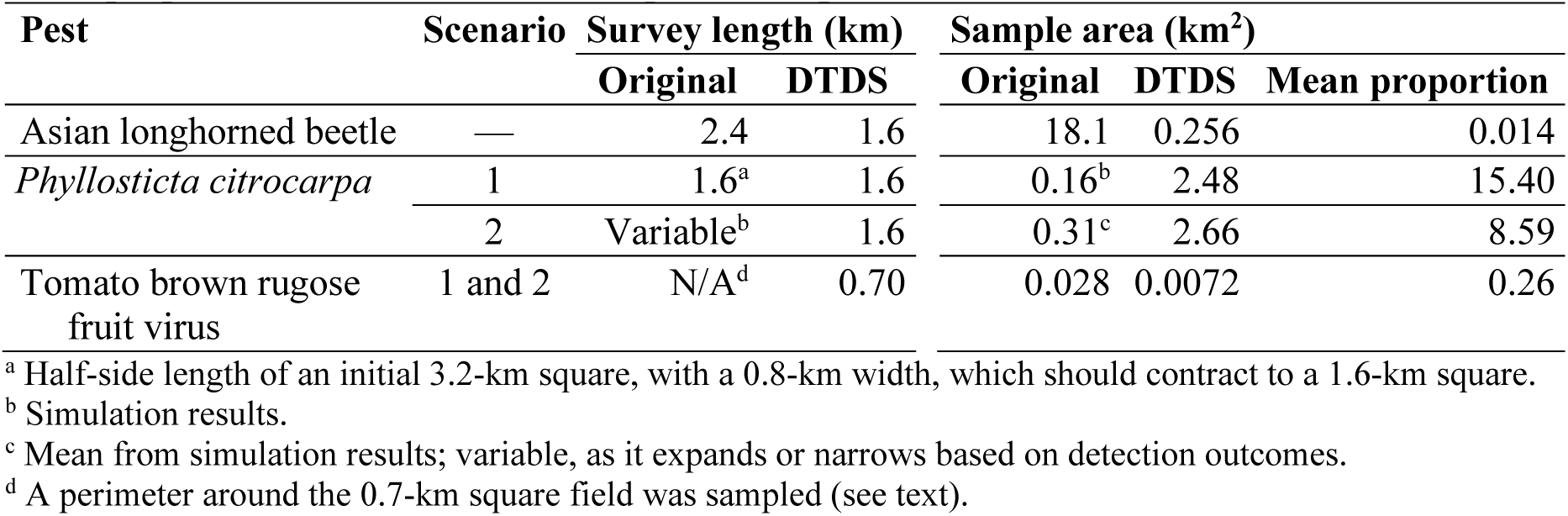
Delimitation survey radii and areas for published and recommended DTDS plans, and mean proportion of DTDS area to the published plan area.

| Pest | Scenario | Survey length (km) |  | Sample area (km<sup>2</sup>) |  |  |
| --- | --- | --- | --- | --- | --- | --- |
|  |  | Original | DTDS | Original | DTDS | Mean proportion |
| Asian longhorned beetle | — | 2.4 | 1.6 | 18.1 | 0.256 | 0.014 |
| *Phyllosticta citricarpa* | 1 | 1.6<sup>a</sup> | 1.6 | 0.16<sup>b</sup> | 2.48 | 15.40 |
|  | 2 | Variable<sup>b</sup> | 1.6 | 0.31<sup>c</sup> | 2.66 | 8.59 |
| Tomato brown rugose fruit virus | 1 and 2 | N/A<sup>d</sup> | 0.70 | 0.028 | 0.0072 | 0.26 |

<sup>a</sup> Half-side length of an initial 3.2-km square, with a 0.8-km width, which should contract to a 1.6-km square.
<sup>b</sup> Simulation results.
<sup>c</sup> Mean from simulation results; variable, as it expands or narrows based on detection outcomes.
<sup>d</sup> A perimeter around the 0.7-km square field was sampled (see text).

##### Test conditions

We set the infested area radius to 2 km, which gave an average of about 3,140 infested trees based on density and the infestation rate.

##### DTDS survey design

ALB is not highly motile (Favaro et al. 2015). Given that, we estimated (based upon Caton et al. 2021) an optimal survey radius of 1.6 km (Table 1). We aimed to detect an infestation rate of 0.01 per tree, which gave mean infested host tree density of 250 per km^2^. At that density, *A*_crit_ = 0.24 km^2^ (equation 1, *X* = 60). That gave *W*_0_ = 0.075 km (equation 3), which was much too large for two surveyors. We therefore used multiple transects, which is common (e.g., Buckland et al. 2007). If each inspector covered at least 4 m (i.e., *W* = 0.008 km), the number of transects (randomly arrayed) needed was 10 (equation 4), which gave total *A* of 0.256 km^2^ (Table 1).

#### Phyllosticta citricarpa

##### Published plan description

The pathogen *P. citricarpa* causes black spot disease in citrus. We based this on the published delimitation survey example for a single epicenter, which specified a host density of 45,000 trees per km^2^ and an infection rate of concern of 0.001 (EFSA et al. 2020b). The example assumes two years since introduction, giving an initial square “potentially infested zone” with sides of 3.2 km (Fig. 3). The outermost 800-m-wide band (Band 1) is inspected first. If no positives are found, the survey continues in the innermost band (Band 0, 1.6 km-square). Alternately, if pests are detected in the first band, the survey *expands* in 800-m increments (i.e., Band 2, Band 3, etc.), stopping only when no more positives are found (Fig. 3). The sampling plan was hypergeometric, with confidence (*C*) = 0.95, infestation rate of concern (*p*_Test_) = 0.001, and sensitivity = 0.8 (EFSA et al. 2020b). This gave sample sizes (*n*) from 3683 to 3737 in each band based on host numbers in each area (see below).

**Figure 3.**
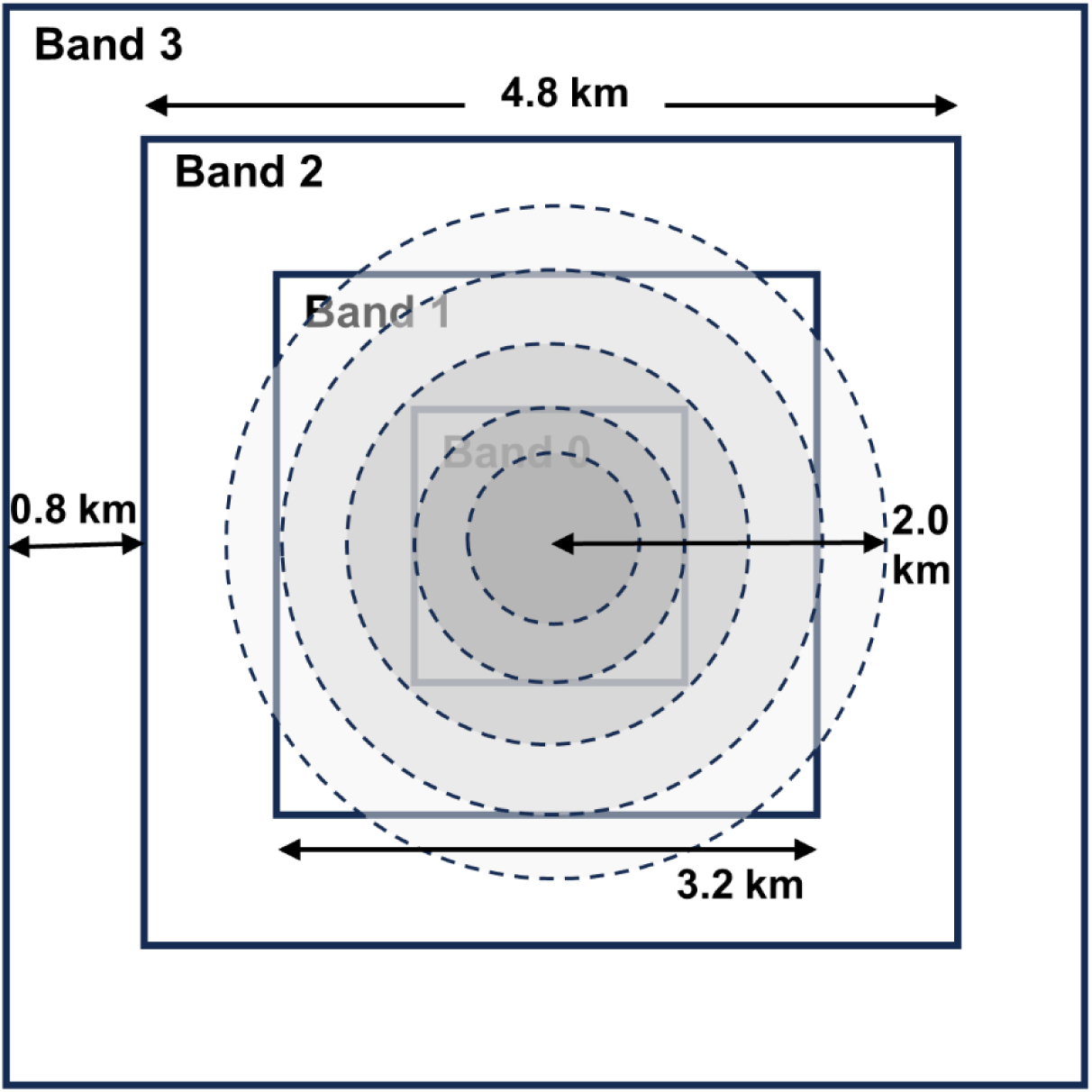
A diagram of the *Phyllosticta citrocarpa* delimitation survey in Scenario 2, in which the infested area extends into Band 2 and has an infestation rate that declines from the epicenter outward (shaded circles). The innermost band, Band 0 (faint), is surveyed if no positives occur in Band 1, while Bands 2 and 3 are surveyed if positives occur in the preceding band. A successful delimitation under the published plan will survey each of Bands 1-3 and skip Band 0.

##### Test conditions

We simulated two scenarios that corresponded to the authors’ examples of infestations that were smaller or larger than the original area of potential infestation. In the <u>first</u> <u>scenario</u>, the spatial extent of the incursion had a 0.8 km radius, requiring a (unsuccessful) survey in Band 1, followed by surveying Band 0. Note that the square Band 0 was slightly larger than the circular infested area, which decreased the effective selection rate somewhat. The infested area had 90,478 trees on average, giving about 90 infested trees with *p*_Inf_ = 0.001. Band 0 had 115,200 trees on average, though, so the selection likelihood (*p*_Select_) was only 0.000781 on average. Given *n* = 3702, the expected mean number of infested trees detected was only 2.9.

In the <u>second scenario</u>, the infested area radius was 2.0 km, extending into Band 2 (Fig. 3). After imposing the square sampling areas from the original plan onto the infested area, we calculated that Band 0 contained 0.444 of the expected total infested trees, Band 1 contained 0.441, and Band 2 contained only 0.115. This directly impacted the likelihood of detecting the infestation in each band. For example, with *p*_Inf_ = 0.01, the mean number of infested trees was 5651, and only 650 were in Band 2. Band 2 had 691,200 trees on average (not shown), giving an effective selection rate of only 0.00094. With a sample size of 3737 the mean number selected was only 3.5, which projected to a substantial failure rate.

##### DTDS survey design

The pathogen is a weak disperser (Perryman et al. 2014). We used a transect radius of 1.6 km in both scenarios, as this is the minimum recommended distance suggested for general delimitation surveying based on pest dispersal capabilities (Caton et al. 2021). Based on the tree density and infestation rate of concern, *m* = 45 infected trees per km^2^, and *A*_crit_ = 1.33 km^2^ (equation 1, *X* = 60). In both scenarios, based on *R* and *m*, the values of *W*_0_ were not practicable. We used additional transects (see below), which equated to adding orchard rows, to achieve *W*_T_ of about 4.7 m (i.e., 1 orchard row) for one inspector. In Scenario 1 we centered transects on the epicenter. To make the DTDS plans more like the published plan in Scenario 2, we sampled the entire area of Bands 1 and 0 (3.2-km square) with parallel, stratified random transects. Random rows were chosen within blocks sized to give the required number of total transects.

#### Tomato brown rugose fruit virus (TBRFV)

##### Published plan description

In this survey plant material will be collected for detection of TBRFV in a lab. An average tomato farm size in California, for example, is about 0.4 km^2^ (NASS 2019). We simulated a 700-m by 700-m field (area = 0.49 km^2^). A normal plant density for tomatoes is 1.2 plants per m^2^, or about 1.1 plants every m, with about 636 rows in 700 m on average. The original delimitation plan in agricultural fields was for area sampling of infected fields, but only in a perimeter at the edge (PPQ 2020b). The perimeter areas are arbitrarily divided into cells (survey units) and a requisite number of hypergeometric samples taken from each cell to give 95 percent confidence in detecting *p*_Inf_ = 0.01. Using 10-by-10 m cells (area = 100 m^2^), the perimeter of the 700-m-square field has 276 cells, with a total area of 27,600 m^2^ (Table 1). Each cell has about 120 plants (*N*). Hypergeometric sampling (Fosgate 2009) gives *n* = 114 per cell, or nearly every plant. The total *n* was 31,464. Note that survey costs would include the lab testing of every sample.

##### Test conditions

As the original plan was field-centric, only the infested field described above was simulated. The infestation was assumed to potentially be present throughout the entire field, with overall *p*_Inf_ = 0.01. We tested both survey plans in two scenarios with differing placements of the epicenter. In scenario 1, the epicenter was in the center of the field (Supp. Fig. S1a), while it was near the lower left corner in scenario 2, 50 m in on the diagonal (Supp. Fig. S1b). In scenario 2 the field radius of 495 m (distance to corner of square) is not the best description of the possible boundary, because the farthest detection could be up to 940 m.

##### DTDS survey design

In this design each row is a potential transect, so we selected from the 636 rows to survey. Each plant along the row would be sampled. At a 1 percent infestation rate, we need to sample 6,000 plants to generate 60 positives. This means we need to sample, on average, about 10 transects (= 6000 / 636; similar to equation 1), with slight dependence on the different scenarios because of epicenter placement.

### Simulation specifications

#### Survey model specifications

Models were all coded in spreadsheets and run using @Risk ver. 7.5.1 Professional Edition (Palisade Corporation, 31 Decker Road, Newfield, NY 14867), a Microsoft Excel add-in. Unless otherwise specified below, simulation settings were as follows: number of iterations = 100,000; sampling type = Latin Hypercube; and random seed = 101.

The basic model structure was to determine the infested area size and the number of infested/infected hosts (positives), and the survey area and potential number of sampled hosts (Fig. 4). In some of the published plans (*P. citrocarpa* and TBRFV) the sample size was modified by subsampling. The ratio of positives in the survey area to the number of potential samples determined the likelihood of selection, which was used to predict the number of detected positives. That result determined the survey outcome. Other metrics were calculated from sample areas and numbers.

**Figure 4.**
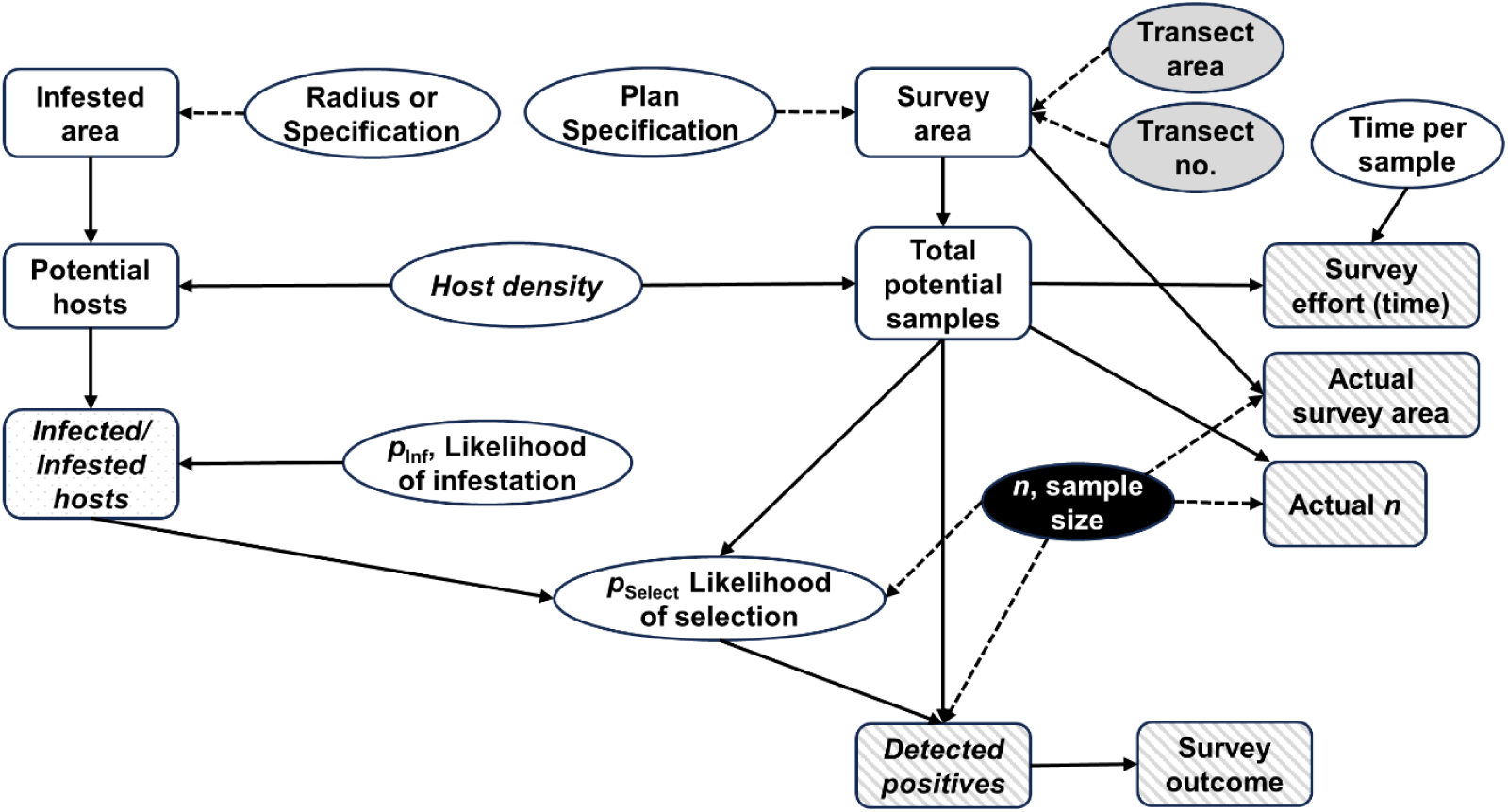
Diagram of the basic simulation model structure for evaluating delimitation survey outcomes based on survey plans. Rectangles are state variables, ovals are parameters or input specifications, and lines show direct (solid) or informational (dashed) dependencies. Gray-shaded ovals represent the transect plan specifications. Cross-hatched variables are outputs, and elements with italicized text are stochastic. The solid black oval and dashed lines indicate specified hypergeometric sample sizes for the published plans for *Phyllosticta citropcarpa* and tomato brown rugose fruit virus.

Parameters were defined directly from the source or were standardized by case study with realistic values (e.g., infestation rates, inspection times) (Table 2). For single (mean) estimate parameters (e.g., plants per km^2^), we always added a moderate amount of uncertainty by using lower and upper values that were ten percent different from the mean.

**Table 2.**
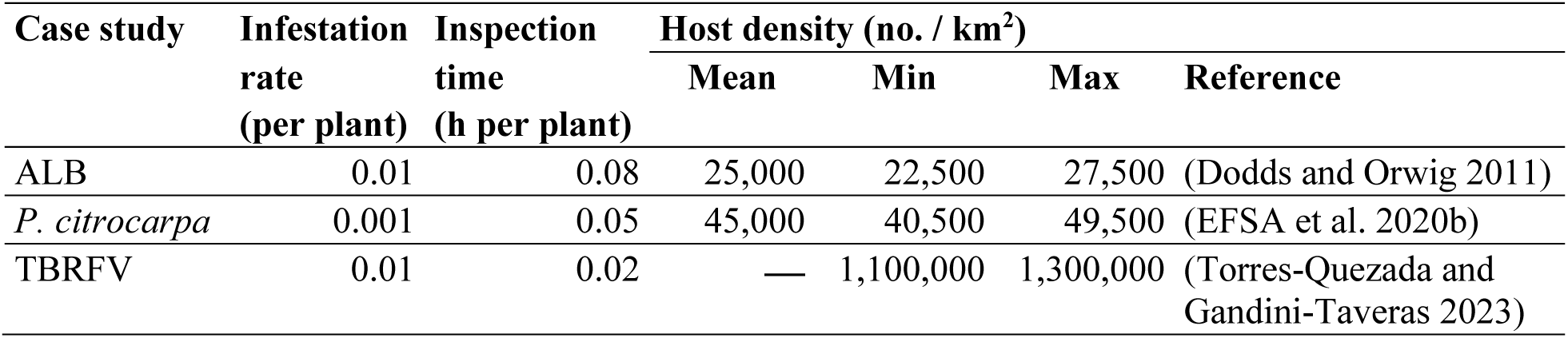
Survey model parameters for three case studies for infestation rates, inspection times, and host densities (means, minima and maxima) with density references.

Many functions were basic arithmetic, such as equations 1-4, as well as area sizes (π × *R*^2^) and infestation densities (no. per unit area). The likelihood of selecting an infected host (*p*_Select_) was the ratio of the number of infected hosts to the number of hosts sampled. Other standard functions were logical, such as survey success (1) or failure (0) based on the numbers of positives found.

The binomial distribution was used extensively for stochastic sampling processes. In that process *N* independent, identical trials are run, each one with the same probability of success, *p*<u>_Success_</u>, producing some number of successes, *s* (Vose 2000):

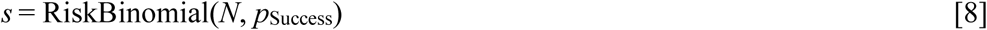

where *N* is the total number of items. For example, this function was used to find the number of positives detected, using *p*_Select_ and *N* equal to *n* (sample size). We assumed perfect detection (i.e., sensitivity = 1.0) across all simulations for simplicity.

We randomly determined host plant densities, *D_s_*, using a uniform probability distribution (i.e., every value equally likely). That general function was as follows:

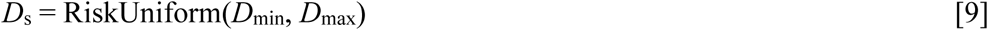

where *D*_min_ is the minimum, and *D*_max_ is the maximum.

#### Spatial model specifications

Completely evaluating the DTDS approach required generating hypothetical distance data for the positives. In most cases we simulated randomly placed transects explicitly to verify that that they found the target number of positives (Fig. 5, detailed model). In one case, for *P. citrocarpa* Scenario 1 (Fig. 5, simple model), we used the defined dispersal distance distribution to randomly sample 65 values (see below). In every case, because locations often were within 3 km of the epicenter, distances from the origin were calculated with the Pythagorean theorem, for simplicity.

**Figure 5.**
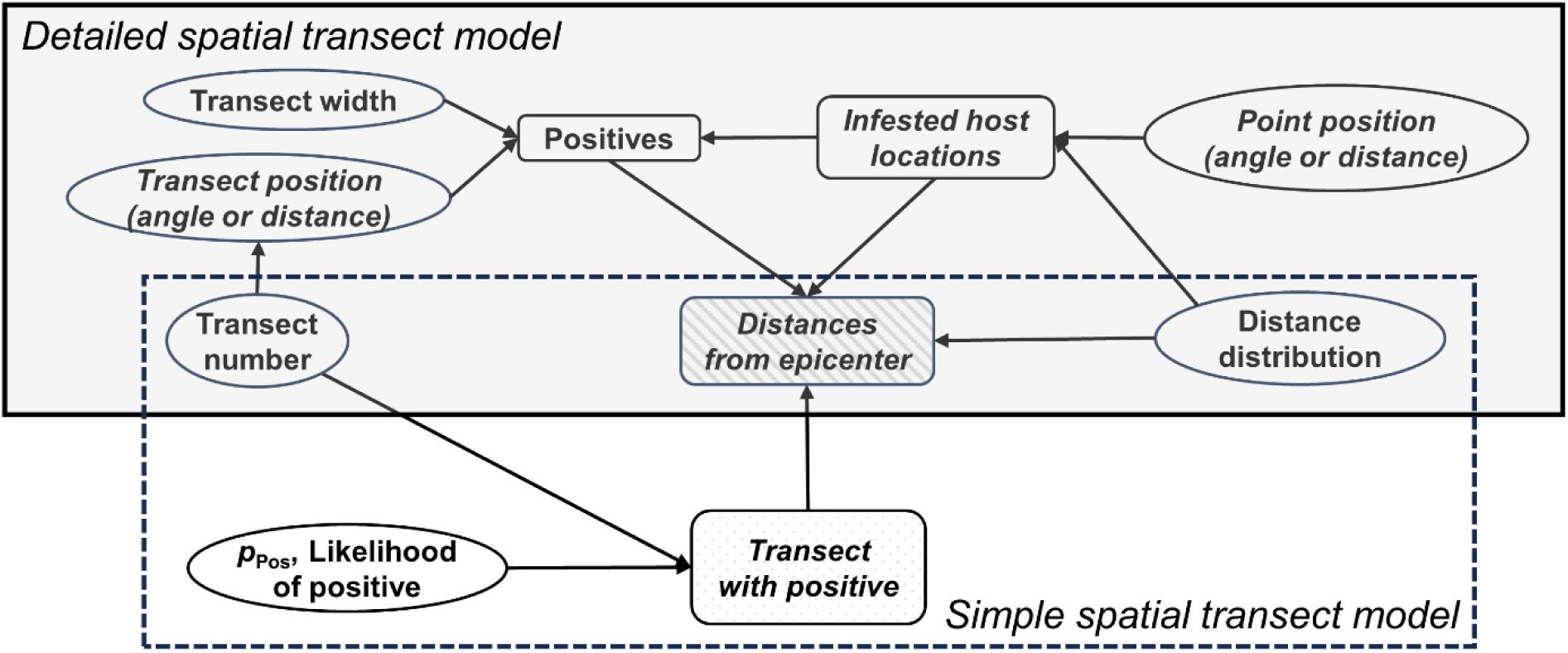
Model diagram for simulating distance data for positives in delimitation surveys. Detailed (shaded box) and simple (dashed box) methods used to generate hypothetical distance data for positives from transect sampling survey plans. Rectangles are state variables, ovals are parameters or input specifications, and solid lines show dependencies. Cross-hatched variables are outputs, while elements with italicized text are stochastic [Note: “Distances from epicenter” is deterministic in the detailed approach.]

##### ALB case study

The locations of 2011 positives (mean from simulations for transect-covered area) were randomly placed within the infested area. Coordinates were projected onto grids with an origin of (0,0) using the following function to find the *X*-axis value (*X*_i_):

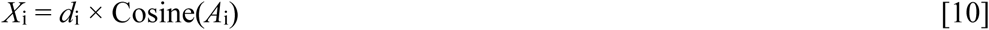

where *d*_i_ is the randomly sampled distance from origin and *A*_i_ is the random angle (equation 9) for *i* positive points. The dispersal density was a linear function with a y*-*intercept of 0.25 and slope =-0.0001, which gave *X* = 0.05 at *Y* = 2000 m. The function for the *Y*-axis value (*Y*_i_) used the same sampled values from equation 9:

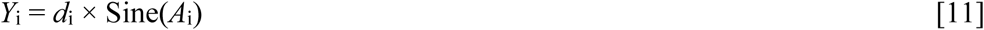

Here we randomly placed 10 transects within the circular survey area, all passing through the epicenter, with a uniform angle (*A*) probability (equation 9). The closest distance from every positive point to each transect was calculated from the associated transect *Y* coordinate based on the hypotenuse length and angle to the transect from each location, using well-known trigonometry functions (not shown). Any location within half the transect width (4.0 m, see above) was a positive.

##### *P*. *citrocarpa* Scenario 1

In this instance we used the simple method (Fig. 3) and randomly selected points from the distance function. That distribution was an exponential decay distribution (equation 5) with *a* = 0.5 and *b* = 0.005, and a boundary of 0.8 km (Supp. Fig. S2). Based on the mean number of detections (below), we used a selection rate over the total number of transects to randomly select transects with positives and then determine the random distance.

##### *P*. *citrocarpa* Scenario 2

In this scenario we randomly placed parallel transects along the side of the square survey area using a stratified scheme. For simplicity, we only simulated the transects that would be placed within the 2.0 km infested area (i.e., could result in a positive), which was 104 of 166 transects. The block width was 38.5 m, which encompassed eight rows. Sampled rows in blocks were chosen with a uniform probability (i.e., a discrete function with integer values from 1 to 8 and equal weights). Values of *p*_Inf_ varied by distance from the epicenter (Supp. Fig. S3) and were chosen to provide approximately the mean number of positives from the survey model (see results below).

##### TBRFV case study

For the published survey, generated positive locations purely for visualization. We randomly located the expected number of detections (from the survey model above) in the perimeter, based on *p*_Inf_ as a function of distance from the epicenter. Values of *p*_Inf_ dropped incrementally from 0.027 to 0.007 every 100 m away from the epicenter (Supp. Fig. S4).

For the DTDS surveys, in both scenarios we randomly selected 10 starting transect locations stratified along one side of the field. The parallel transects extended 300 m across the field. Positives occurred randomly in 1-m segments with *p*_Inf_ varying by distance from the epicenter, as above (Supp. Fig. S4).

### Evaluating curve fitting methodologies

#### Analysis

The standard analyses used the baseline (unaltered) data. For EVA POT, we defined the threshold based on the “mrlplot” and “threshrange.plot” charts in the R package extRemes (2.0), which used the raw distance data. Suggested threshold values were tested by excluding data below the threshold and then verifying that the resulting histograms seemed appropriate (i.e., skew and shape).

For curve fitting, data were converted to histograms of the proportions of detections as a function of distance from the epicenter. We calculated the ideal number of bins in the histogram according to Sturge’s Rule (Scott 2009), which is typically useful for *n* < 200. Histogram data, using midpoints of the intervals, were fit in JMP (ver. 13.1.0; RRID:SCR_014242) to the nonlinear, two-parameter exponential model (equation 5). To evaluate fit we compared the Akaike Information Criteria (Hilborn and Mangel 1997) results for a three-parameter exponential decay model, as follows:

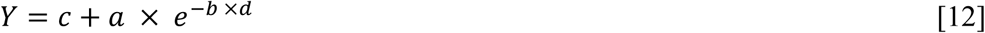

where *c* is a lower limit parameter. For *c* = 0, equation 12 becomes equation 5.

#### Distribution variability

For each case study and scenario above we compared curve fitting and boundary setting results from fitting the baseline data to fitting the data in the EVA-POT technique.

#### Clustering

In the case studies, the distributions of positives usually varied as a function of distance from the epicenter but were otherwise homogeneous. To demonstrate that our novel approach could handle clustered populations (Leung et al. 2010), here we examined the ability of transects to cover non-homogeneous distributions of positives. We created four grids of random locations that varied from relatively homogeneous to highly clustered, using a Thomas point process (Baddeley et al. 2015). We did this in the R package spatstat.random, using the function “rThomas.” The function has three parameters: as follows:

- Scale: the standard deviation of random displacement (along each coordinate axis) of a point from its cluster center
- μ (mu): the mean number of points per cluster
- κ (kappa), the intensity of the cluster centers

We used constant scale = 0.15, while varying μ from 1 (Pattern 1), to 6 (Pattern 2), 12 (Pattern 3), and 24 (Pattern 4). To generate the expected declining distributions with distance, we specified κ as an exponential function (equation 5), with a distribution that had either low or high skewing (Supp. Figs. 5 and 6). For unknown reasons, the process produced fewer points near the epicenter than needed, so we increased the counts of the first two bins (100 m width) by factors of 8 and 4, respectively for all datasets. We randomly sampled 60 points from those distributions in @Risk for testing.

## Results

### Curve fitting and boundary setting with clustering

Fitting the data via the standard approach (all data) always provided reasonable fits but for the more skewed distributions it failed in every case to provide a sufficient 99^th^-percentile (radial) boundary estimate (Table 3). The curve fits from EVA-POT were similarly reasonable and 99^th^-percentile (radial) boundary estimates were adequate in every case, although sometimes well beyond the actual infested radius.

**Table 3.** Exponential curve fitting results with either the standard method (full data) or the extreme values analysis peaks-over-threshold (EVA POT) method for hypothetical presence data with less or more skewing and four levels of clustering (see text). Metrics shown are scale and rate parameter values, root mean squared error (RMSE), R^2^ value, the actual extent and estimated boundary (99^th^ radial percentile), and outcome.

| Method | Skewness | Clustering | Scale | Rate | RMSE | $R^2$ | Boundary (m) | | Outcome |
| --- | --- | --- | --- | --- | --- | --- | --- | --- | --- |
|  |  |  |  |  |  |  | Actual | Estimated |  |
| Standard | Less | 1 | 0.5271 | -0.0019 | 0.0348 | 0.951 | 2,084.3 | 3,540.5 | Success |
| Standard | Less | 2 | 0.5998 | -0.0023 | 0.0177 | 0.988 | 2,053.8 | 2,912.8 | Success |
| Standard | Less | 3 | 0.4221 | -0.0013 | 0.0311 | 0.941 | 2,080.7 | 5,036.3 | Success |
| Standard | Less | 4 | 0.6461 | -0.0021 | 0.0378 | 0.960 | 2,188.6 | 3,206.9 | Success |
| Standard | More | 1 | 1.1340 | -0.0039 | 0.0236 | 0.993 | 1,971.2 | 1,960.0 | Failure |
| Standard | More | 2 | 1.1458 | -0.0038 | 0.0219 | 0.993 | 2,115.2 | 1,739.2 | Failure |
| Standard | More | 3 | 0.9929 | -0.0035 | 0.0454 | 0.966 | 1,998.6 | 1,920.8 | Failure |
| Standard | More | 4 | 0.8753 | -0.0040 | 0.0570 | 0.921 | 1,869.7 | 1,680.6 | Failure |
| EVA POT | Less | 1 | 3.7292 | -0.0027 | 0.0192 | 0.995 | 2,084.3 | 2,472.4 | Success |
| EVA POT | Less | 2 | 0.9208 | -0.0019 | 0.0416 | 0.934 | 2,053.8 | 3,446.7 | Success |
| EVA POT | Less | 3 | 0.4303 | -0.0007 | 0.0584 | 0.632 | 2,080.7 | 9,416.1 | Success |
| EVA POT | Less | 4 | 0.8353 | -0.0019 | 0.0702 | 0.892 | 2,188.6 | 3,471.9 | Success |
| EVA POT | More | 1 | 1.0240 | -0.0018 | 0.0653 | 0.928 | 1,971.2 | 3,712.7 | Success |
| EVA POT | More | 2 | 0.9807 | -0.0021 | 0.0649 | 0.899 | 2,115.2 | 3,206.9 | Success |
| EVA POT | More | 3 | 1.0317 | -0.0023 | 0.0655 | 0.882 | 1,998.6 | 2,936.0 | Success |
| EVA POT | More | 4 | 0.6181 | -0.0014 | 0.0261 | 0.969 | 1,869.7 | 4,745.1 | Success |

### Case studies: Boundary Setting and Outcomes

#### Asian longhorned beetle (ALB)

The mean number of infested trees detected in the original plan was 3141.6 (Fig. 6a), compared to 64.0 in the DTDS design (range = 32 to 107; Fig. 6b). The DTDS survey reduced *n* by almost 99 percent (Table 4). The estimated time to complete the survey decreased by the same proportion, saving 4619 person-days.

**Figure 6.**
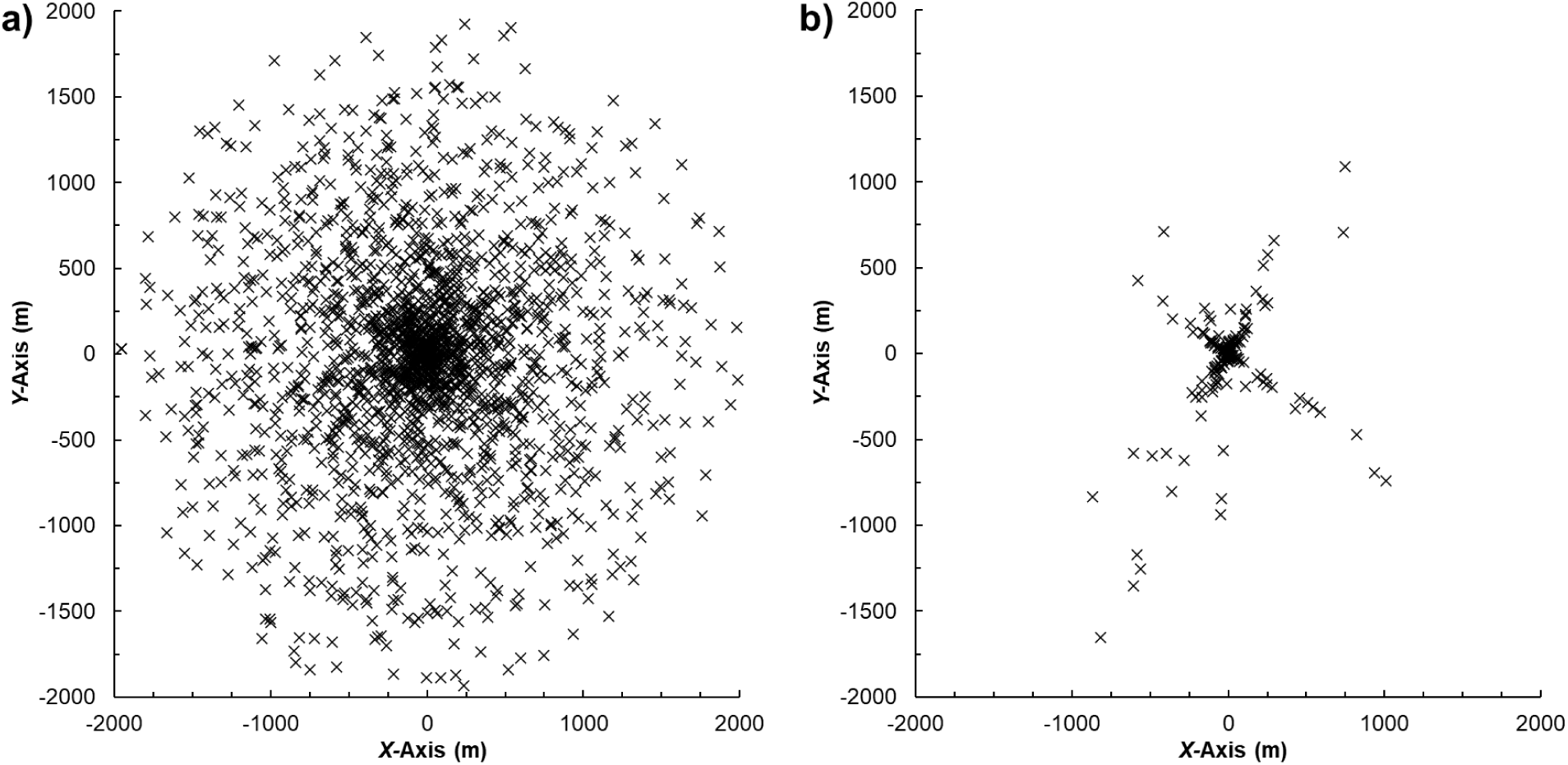
Simulated ALB delimitation survey detections. Example simulated delimitation field survey results for two designs for ALB, showing (a) positives in the original plan, and (b) a transect-based plan (DTDS) designed to give around 60 positives, but which actually produced 174 detections in the stochastic exercise.

**Table 4.**
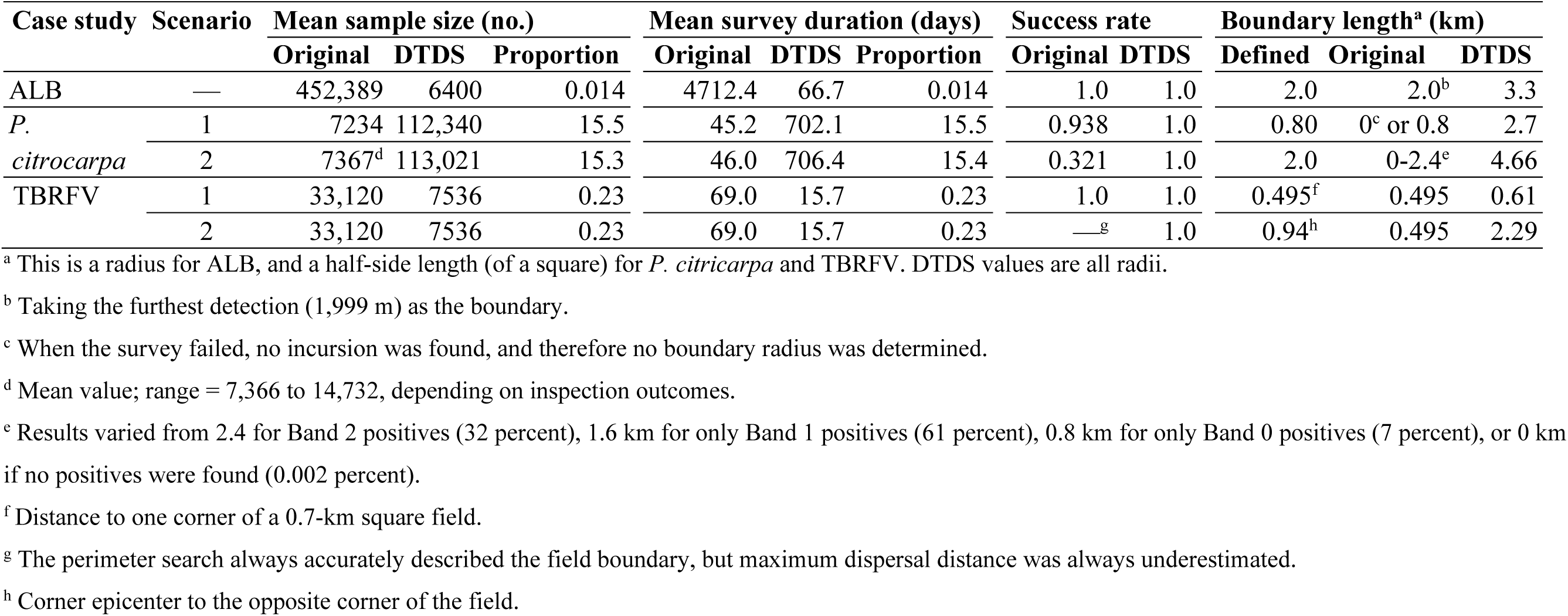
Simulation results for observation-based delimitation surveys in three case studies, for Asian long-horned beetle (ALB), *Phyllosticta citrocarpa*, and tomato brown rugose fruit virus (TBRFV). Sample areas and sizes for localized observation-based delimitation surveys in simulations of three case studies for both the original and DTDS survey plans, showing proportions for the DTDS surveys relative to the original plans.

The threshold for the EVA POT curve fitting was 500 m. The 99^th^ percentile radial estimate of the boundary distance from the fitted curve was 3.3 km. This overestimated the test boundary by 66 percent (Table 4; Fig. 7). In contrast, the boundary from the same percentile for the baseline function was only 1004 m, an underestimation by 50 percent. For reference, a radial percentile of 91 gave a boundary just beyond the test distance.

**Figure 7.**
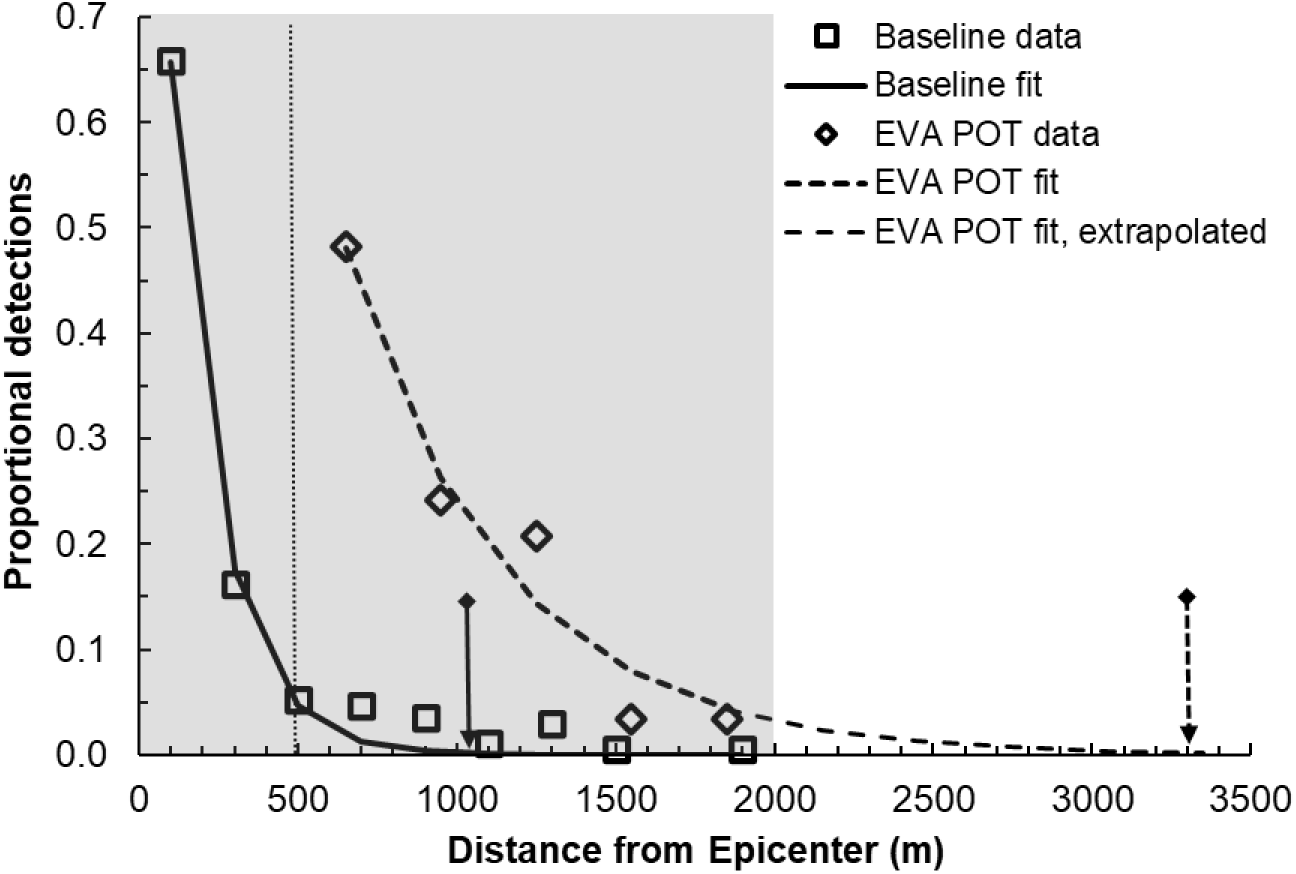
Simulated and fitted detection distances for ALB. Proportional detections as a function of distance from the origin for a simulated delimitation survey data for ALB, and fitted exponential decay functions to either the full (baseline) data or the extreme values analysis peak-over-threshold (EVA POT) data, with an extrapolation for the EVA POT curve fit. The dotted vertical line shows the EVA POT threshold, and the arrows show the 99^th^ radial percentiles for the baseline (solid line; 1004 m) or the EVA POT data (dashed line; 3303 m).

#### Phyllosticta citricarpa

##### Scenario 1

In the EFSA survey plan the mean number of detections was only 2.8, and the survey failed to detect *any* positives about 6 percent of the time, implying no incursion at all. That value would increase if a more realistic inspection sensitivity were used.

The DTDS survey with 165 transects never failed to detect positives, finding an average of 56.1 infected hosts per survey (range = 35 to 76). But *n* was more than 15 times greater (Table 4). For reference, a hypergeometric sample that would give *n* approximately equal to the mean trees sampled in the DTDS plan would require decreasing the infestation level from 0.001 to 0.00003 (it’s not possible to only raise the confidence level and reach the required mean *n*).

For this scenario the EVA POT threshold was 300 m. The 99^th^ radial percentile from the fitted curve was 2.7 km (Table 4; Fig. 8), which overestimated the test boundary by 233 percent. This would give a potential management area of 22.9 km^2^. Although this was ten times larger than the actual infestation here, it robustly accounted for uncertainty. For reference, the 59^th^ radial percentile gave a boundary near to the actual boundary of 788 m.

**Figure 8.**
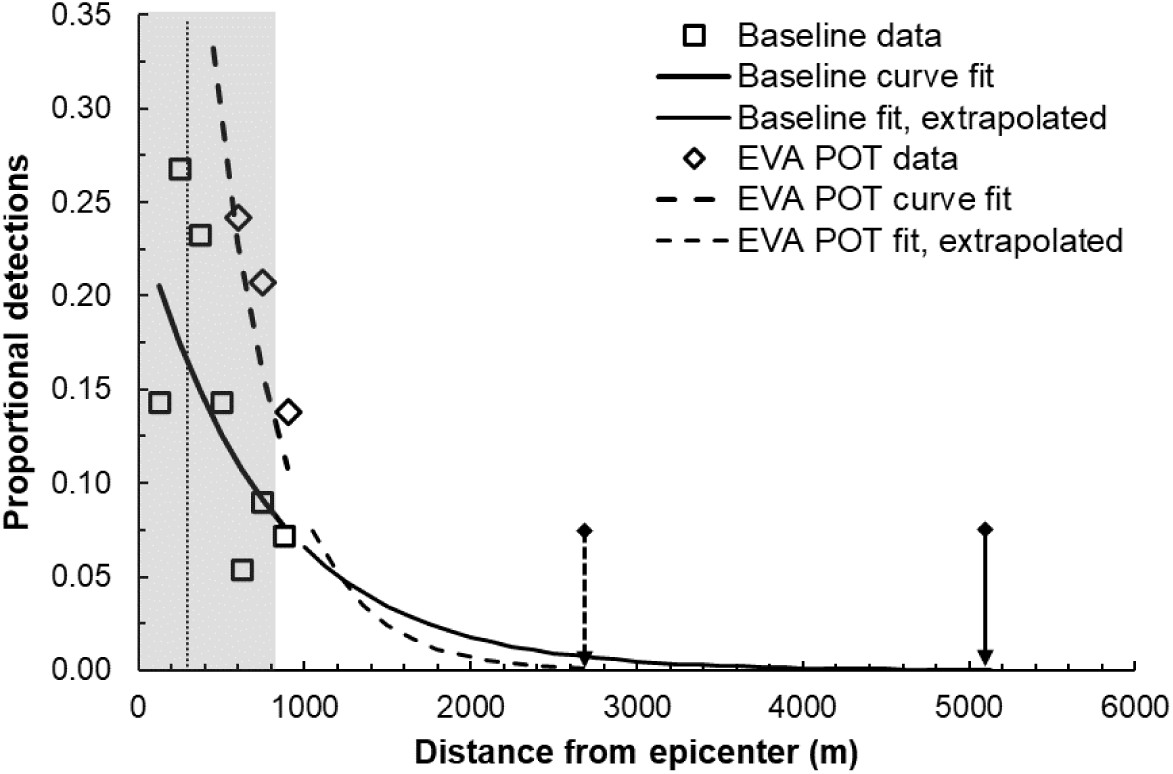
Simulated and fitted detection distances for *Phyllosticta citricarpa* Scenario 1. Proportional detections (symbols) as a function of distance from the epicenter for simulated delimitation survey data for *P. citricarpa*, and fitted exponential decay functions (thick lines) to either the full (baseline) data or the extreme values analysis peak-over-threshold (EVA POT) data, with extrapolations (thin lines). The shaded area is the actual area of extent, the dotted vertical line shows the EVA POT threshold, and the arrows show the 99^th^ radial percentiles for the baseline (solid line; 5091 m) and the EVA POT data (dashed line; 2661 m). [Histogram binning accounts for points that seem to be outside the infested area.]

##### Scenario 2

Here the original plan had a minimum and mean sample size of 7367 trees (Table 4), but the maximum was 14,732 depending on outcomes (narrowing or expanding). However, the survey failed to detect infected trees in Band 1 almost 7 percent of the time, leading in those cases to a *narrowing* of the infested area (Table 4). Even when correctly surveying Band 2, detection there failed in 66 percent of iterations, because infested trees were sparse. Finally, infested trees in Band 0 were missed 0.002 percent of the time, giving no detection of the incursion at all. The overall success rate was therefore only 32.1 percent (Table 4).

By contrast, the DTDS plan required sampling 166 transects, giving a much larger *n* than the original plan (Table 4), and a mean detection number of 61.6 infested trees (range = 28 to 106), as planned. The EVA POT threshold was 300 m, and the 99^th^ percentile radial boundary estimate was 4.66 km (Table 4; Fig. 9). This overestimated the test boundary by 133 percent and gave a potential management area of 62 km^2^, but the survey failure rate was zero, which we think justified the increased survey duration. For reference, the 77^th^ radial percentile gave a boundary near to the actual boundary of 1933 m.

**Figure 9.**
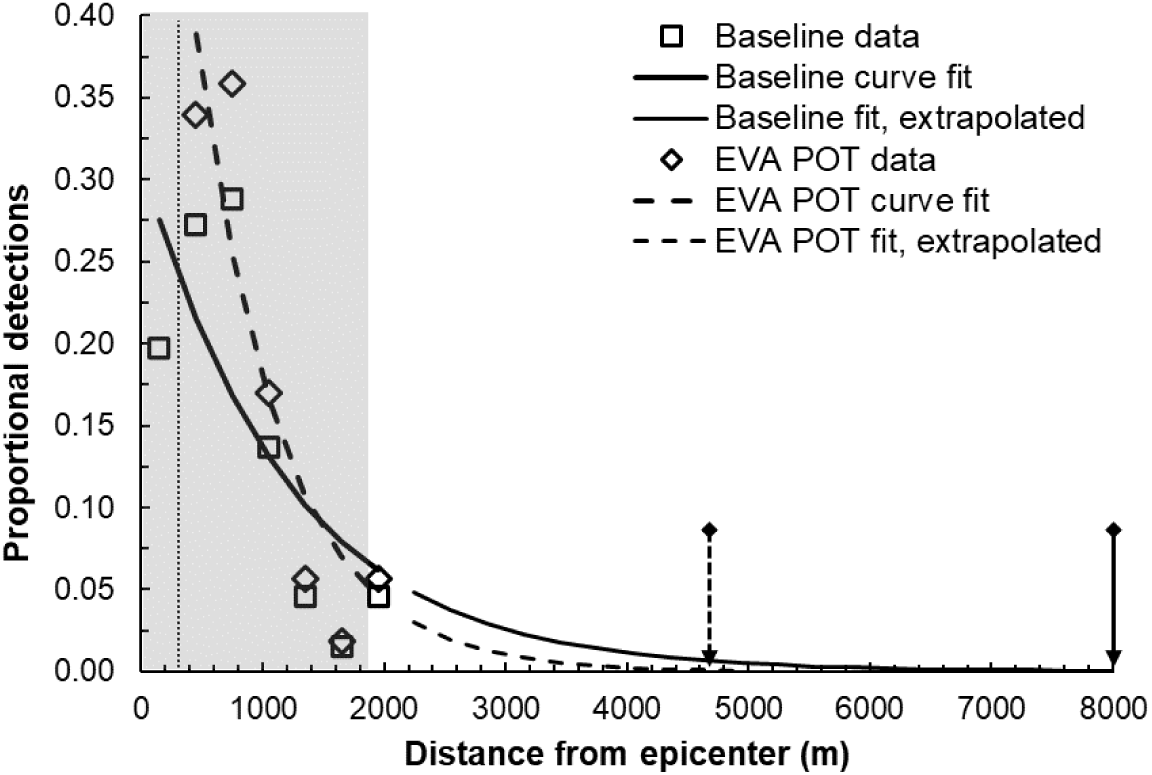
Simulated and fitted detection distances for *Phyllosticta citricarpa* Scenario 2. Proportional detections (symbols) as a function of distance from the origin for simulated delimitation survey data for *P. citricarpa*, and fitted exponential functions (thick lines) to either the full (baseline) data or the extreme values analysis peak-over-threshold (EVA POT) data, with extrapolations (thin lines). The shaded area is the actual area of extent, the dotted vertical line shows the EVA POT threshold, and arrows show the 99^th^ radial percentiles for the baseline (solid line; 8017 m) or the EVA POT data (dashed line; 4655 m). [Histogram binning accounts for points that seem to be outside the infested area.]

#### Tomato brown rugose fruit virus (TBRFV)

##### Scenario 1

The original plan detected a mean of 223.5 positives in the perimeter, while the DTDS plan detected 75.4 plants on average (Table 4). The sample area in the DTDS plan was about 73 percent smaller and *n* was 77 percent lower. Thus, the time required decreased by 77 percent. Those savings would accumulate for every additional field surveyed. Moreover, the perimeter survey gave little information about the potential for spread beyond this single field. With DTDS the EVA POT technique had a threshold of 250 m and the 99^th^ radial percentile boundary radius was 0.61 km (Table 4; Fig. 10). This was 23 percent longer than the actual extent but provided useful information for extending surveys to nearby fields. For reference, the 97^th^ radial percentile gave a boundary estimate very near to the actual extent of 495 m.

**Figure 10.**
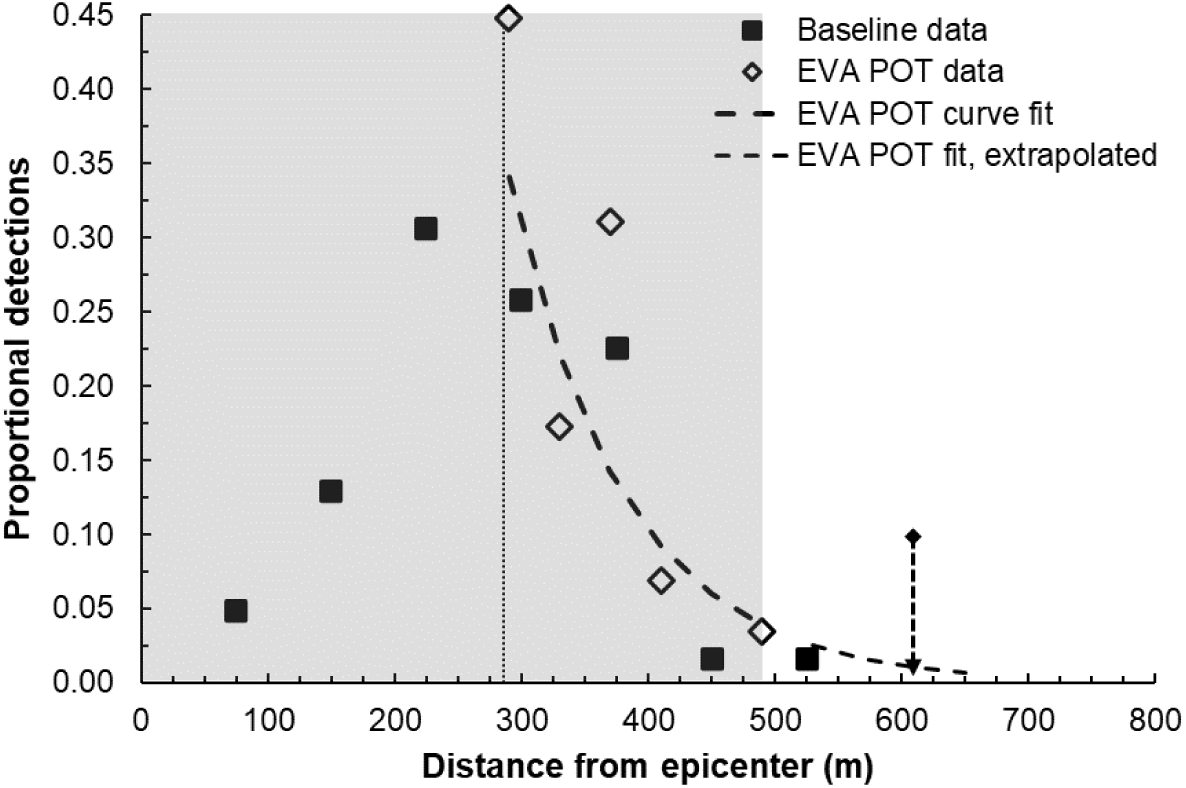
Simulated detection distances for TBRFV in Scenario 1. Proportional detections (symbols) as a function of distance from the origin for simulated delimitation survey data for TBRFV and the fitted exponential function (thick line) to the extreme values analysis peak-over-threshold (EVA POT) data, with an extrapolation (thin line). The full (baseline) data was not amenable to fitting. The shaded area is the actual area of extent, the dotted vertical line shows the EVA POT threshold, and arrows show the 99^th^ radial percentiles for the EVA POT data (dashed line; 610 m). [Histogram binning accounts for points that seem to be outside the area of extent.]

##### Scenario 2

The original plan detected a mean of 342.6 positives in the perimeter (Table 4, Fig. 11a). This increased from the earlier scenario because the infestation rate in the perimeter was much greater (Supp. Fig. S1b). Note that unless the data were inspected carefully, the offset epicenter might not have been identified. The DTDS plan detected 69.3 plants on average, with 60 percent smaller *n* (Table 4; Fig. 11b). Thus, survey duration decreased by almost 78 percent, as above.

**Figure 11.**
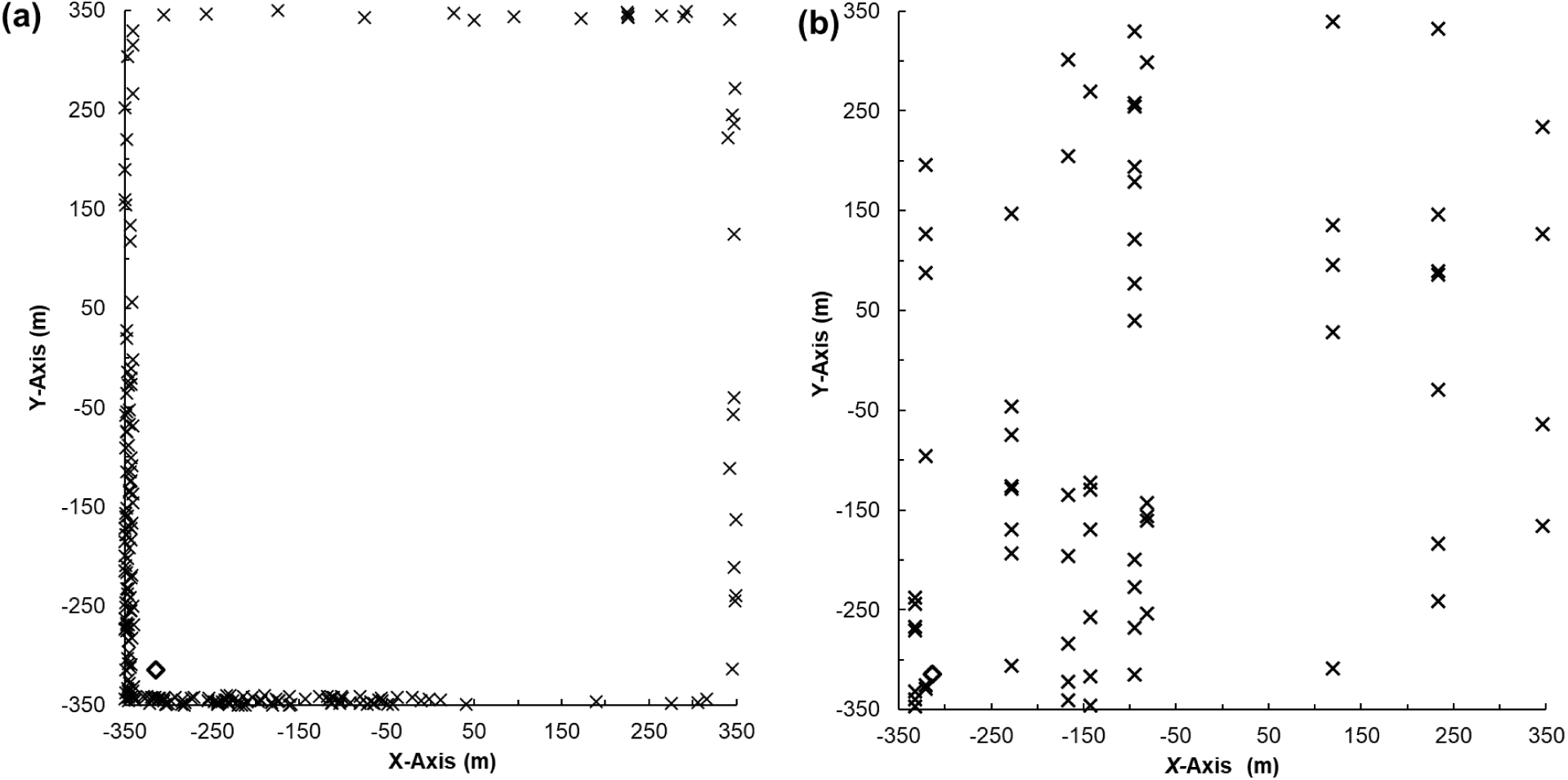
Simulated delimitation survey results for TBRFV for Scenario 2. Simulated delimitation survey results for two designs for TBRFV, with an off-center epicenter (empty diamond). a) The original plan with perimeter sampling of a 10 m-wide area, and B) a DTDS plan, which had stratified randomly placed transects along the *X-*axis. See text for more details.

Assuming the epicenter was in the center of the field yielded a problematic distance distribution (Fig. 12a). The mean value for the *X* coordinate was -90.8 and the mean value for the *Y* coordinate was -65.7, indicating the true epicenter was toward the lower left (Fig. 11b). We tested epicenters with those values, and then in 50-m increments starting at (-100,-100) by recalculating distances and histograms. An epicenter at (-150,-150) gave an acceptable distribution, and with a threshold of 200 m the EVA POT curve fitting yielded a boundary radius of 2.29 km (Table 4; Fig. 12b). This was 144 percent longer than the test radius. For reference, the 76^th^ percentile gave a boundary estimate very near to the actual extent of 940 m.

**Figure 12.**
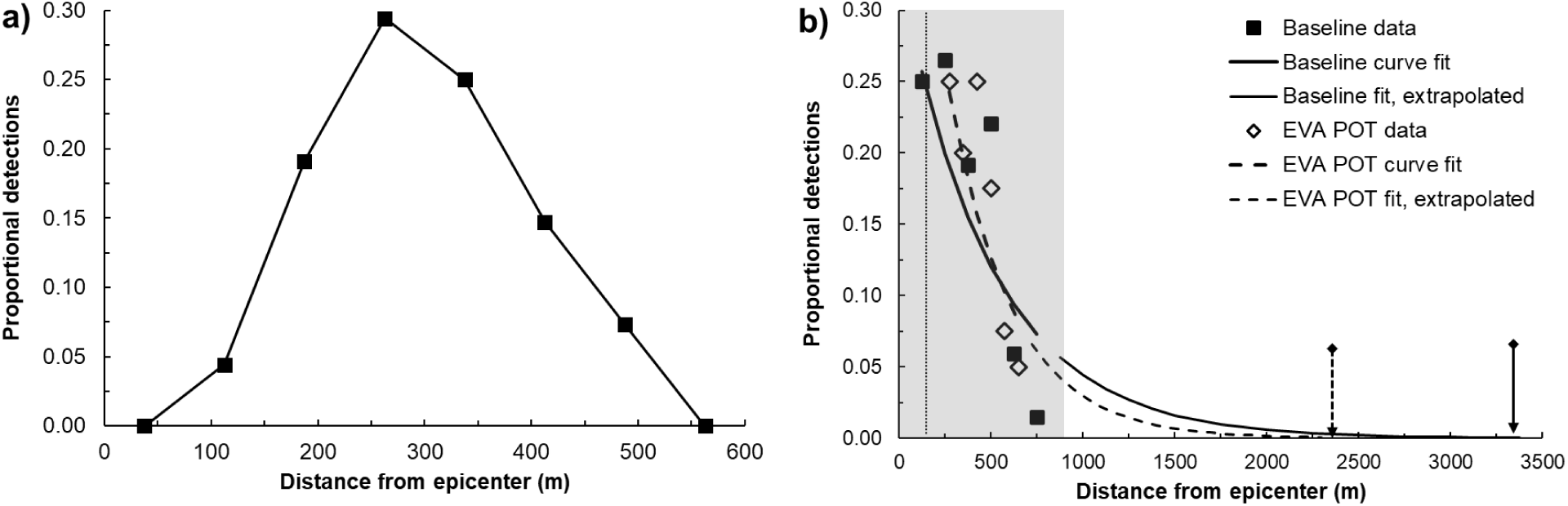
Simulated probability densities for TBRFV in Scenario 1. Simulated probability densities (symbols) of TBRFV as a function of distance from the epicenter for the DTDS survey, for a) the epicenter as the middle of the field and b) the epicenter estimated as (-150,-150). Chart (b) shows the fitted exponential function (thick lines) for the full (baseline) data and for the extreme values analysis peak-over-threshold (EVA POT) data, with extrapolations (thin lines). The shaded area is the actual area of extent, the dotted vertical line shows the EVA POT threshold, and arrows show the 99^th^ radial percentiles for the baseline data (solid line; 3300 m) and for EVA POT data (dashed line; 2291 m).

## Discussion

### General findings

Using a survey design process focusing on delimitation facilitates both quick decision making and avoiding unnecessary resource expenditures, especially if eradication will not be tried (Simberloff 2003; Welsh et al. 2021). Set boundaries can be used quickly to create any needed quarantine areas and then to facilitate management planning and operations. DTDS is a straightforward approach to designing customized, relatively rapid observation-based delimitation surveys. Users specify a transect-based design likely to detect the adventive pest about 120 times, run the survey and collect the positives data. They then fit a curve to extremes in the data and estimate the boundary with uncertainty. The survey details—length, width, and number of transects—can vary greatly depending on the situation, but customization is a key feature of DTDS. Transects are relatively easy to implement in the field (Anderson et al. 1979), and using that tactic to generate data for curve fitting and focusing on occupancy should produce effective and efficient surveys (Table 4), especially in comparison to whole-area surveys like that for ALB. Transect-based delimitation surveys are now being used in some agency response plans (e.g., PPQ 2022b), and could be considered by survey managers facing difficult resource constraints, searching for a quicker process, or desiring more certainty in boundary setting for management.

Completing delimitation surveys requires effort and resources. Sometimes a program may need to inspect 110K trees (Table 4) or cover 200 km^2^ (Schroeder 2012). Even the fastest (hypothetical) survey designs based on DTDS here required many days to complete, and those estimates only accounted for the surveying activity itself, not training, preparation, travel, or analysis and reporting. Given the time and resources required, getting an accurate and meaningful result in a shorter time frame is critical. While we think no supremely easy approach to delimitation currently exists, DTDS seems to improve upon previous approaches that were often much less efficient and sometimes subject to failure.

DTDS differs from the earlier ADD proposal of Leung et al. (2010) in at least six ways:

1. Defining a “fixed” transect length based on information about the species and the situation, and complete sampling of the area when practicable
2. Estimating the total survey area as the size needed to produce at least 120 positives (though 60 positives performed adequately in our tests)
3. Sampling the entire survey (transect) area when practicable, or using stratified random sampling of the survey area otherwise
4. Using EVA POT for curve fitting
5. Accounting for radial dispersal of species during boundary estimation
6. Scouting for boundary verification, and as a recommended pre-survey exercise to improve planning and survey design

In two case studies, DTDS designs were much more efficient than the original plans, reducing sample areas and numbers by 77 percent for TBRFV and by 98 percent for ALB (Table 4). When survey effort can be reduced without jeopardizing the outcome, the financial benefits are clear, and getting faster results would likely further reduce pest response costs (Alvarez and Solís 2018).

In the one exception, for *P. citrocarpa*, the DTDS design increased survey effort greatly but eliminated survey failures that existed with the original plan in both scenarios. Using hypergeometric sampling in delimitation surveys was clearly problematic; *C* = 0.95 (EFSA et al. 2020b) *guarantees* an expected failure rate of 5 percent, and other situational issues can increase the likelihood of failure (Table 4; Fig. 3). We also note that the *P. citricarpa* scenario outcomes would be even worse if *p*_Inf_ had been lower than the test value of 0.01. [The TBRFV survey plan avoided this problem by applying hypergeometric sampling *within* cells, rather than to the overall area, but that plan was still not optimal.] The 68 percent failure rate in scenario 2 was alarming because of its magnitude, but principally because the errors are likely to boost management costs (e.g., Panetta and Lawes 2005).

These results have been corroborated in recent studies. Dynamic delimitation surveys using hypergeometric sampling, as in the EFSA guidelines (EFSA et al. 2020a) (but circular), were evaluated in two different studies (Koh et al. 2025; Sun et al. 2025). Surveys in Koh et al. (2025) targeted the pathogen citrus greening, with *C* = 0.95, *p*_Test_ = 0.01. With sensitivity = 1.0 and a radius that matched dispersal, two designs were fully successful at containment about 34 or 48 percent of the time, but most designs had percentages of about 4 percent or less. Results depended on factors such as accuracy of radius specification, clustering, and starting point. Sun et al. (2025) confirmed that these designs sometimes fail to find *any* incursion, and in the best case the containment success rate was only about 15 percent. Both studies demonstrated that setting boundaries based on the furthest positives is problematic precisely because infestations become sparse at the edges (Lawes and Panetta 2004; Leung et al. 2010; Turgeon et al. 2010). Using hypergeometric sampling makes it even more difficult, since failure rates are built in (Sun et al. 2025).

Delimitation survey designs need to minimize the chance of containment failure. We think this may best be accomplished by avoiding the dynamic methods tested above and hypergeometric sampling, and instead use a method that quantitatively sets a boundary with uncertainty. Very few published survey plans have explicitly discussed setting boundaries based on survey results. In many entire areas were surveyed (e.g., Scott and Batchelor 2014; CDFA 2020; PPQ 2020a), which is a resource-intensive activity more appropriate for determining population size than spatial extent. They automatically fail if the chosen area does not completely contain the pest population (Barclay 2021), such as with *Chrysanthemoides monilifera* subsp. *rotundata*. (L.) Norl. [Asteraceae] (Scott and Batchelor 2014). More dispersive pests present greater difficulties in accurately specifying survey bounds (e.g., Turgeon et al. 2015). For large unsurveyed areas of potential establishment the probability is high that at least one undetected occupied site exists (Becker et al. 2022). Thus, whole-area survey results may be ambiguous at best, or overconfident at worst, about the full spatial extent of the pest population (e.g., Jacobs et al. 2014). DTDS, on the other hand, provides direct evidence that containment has been achieved.

Moreover, the DTDS method has three known safeguards against boundary misspecification. This is important because underestimation of the boundary—failed containment—is the primary concern (Mangel et al. 1984) and can be a major contributing factor to discontinued or unsuccessful pest eradication programs (Howell 2012). The first safeguard is in increasing survey efforts if the recommended number of positives is not met. Using transects helps ensure that the additional effort would be manageable. The second is using EVA POT curve fitting and percentile-based radial estimation of the boundary. Analytical approaches in general seem preferable. In the numerous examples above, EVA POT curve fitting always produced boundary estimates which contained the pest population over varying skewness levels, and with varying levels of non-homogeneity. Although the containment areas were often much larger than the actual spatial extent, overestimation is preferable because it avoids the potential for very large management costs in the future to deal with escaped populations (Woldendorp and Bomford 2004; Panetta and Lawes 2005; Harris and Timmins 2009). Using 120 positives instead of 60 could decrease overestimation. Third, DTDS explicitly calls for post-survey scouting to try to verify the outcome. The objective of post-survey scouting is to try to *invalidate* the boundary by finding target pests outside it.

It may be rare for initial detections to be substantially distant from the main population source (Meats et al. 2003), but misspecification of the origin may sometimes happen (Leung et al. 2010; EFSA et al. 2020b). However, that may not need to be corrected before running the survey. A distribution of distances with an obvious hump indicates a problem. Adjusting the origin— perhaps in conjunction with scouting to verify the new epicenter—and recalculating distances should correct the problem.

### Survey design considerations

The key assumption with distance sampling is that the line transects are placed independently of the organism locations (Buckland et al. 2015), which should always be possible with DTDS. Moreover, we do not need to estimate abundance, which has been a focus of performance studies (e.g., Nomani et al. 2012). Occupancy models should meet some other assumptions (e.g., objects directly on the line are detected with certainty; positives are independent) (Guillera-Arroita et al. 2011; MacKenzie et al. 2013; Buckland et al. 2015), but estimating spatial extent in this context does not require invoking those models. Thus, the approach here likely eliminates some technical concerns. If the survey generates enough positives for analysis, “misses” seem less problematic, which is not always true (e.g., Amburgey et al. 2019). Practices to increase the number of positives, such as pre-survey scouting, or using information from aerial photos, or spatial host or habitat data to better locate transects (Ringvall 2007), should be valid and helpful.

The DTDS designs created in the case studies above were flexible regarding the shapes and sizes of survey areas (e.g., Fig. 6 and the ALB case study), which was already a well-known feature of transects (Buckland et al. 2015). Crossed and parallel line designs both can cover a variety of shapes adequately. When multiple transects are needed, stratification can be applied if warranted. If a relatively large number of transects is required, transects ideally should not overlap and the design should minimize the possibility that an organism could be detected from more than one line (Anderson et al. 1979). [Natural movement of animals from one place to another is not a concern.] In those cases, regular spacing, as opposed to random spacing (Buckland et al. 2015), or adjusting dimensions or shapes may limit interference between lines. Notably, random or regular spacing of line transects had little impact on survey performance with randomly distributed or clumped populations (Nomani et al. 2012). Our results demonstrated the same.

One valid concern is a strong density gradient that results in mean *m* poorly representing the target species density in the survey area. Sparse (or nonexistent) populations in significant portions of the survey area could give *s* << 120. This may be unavoidable in some cases, although pre-survey scouting (or a pilot trial) could help. The fix is relatively straightforward, however: add transects or adjust their lengths and continue to survey until *s* = 120. For rare species, the best strategy is sampling more units less intensively (MacKenzie et al. 2013).

### Management implications

The literature about “delimitation” is sometimes ambiguous. Abundance-estimating surveys seem geared more directly toward management. For example, Lazaro et al. (2020) applied a modeling-based approach to “delimiting surveys” for locating pests within a previously demarcated area. Likewise, in a successful eradication program for blackberries, over half of the costs were for “delimitation” (Parkes and Panetta 2009). That activity, however, was described as “extensive systematic searches” near known infestations (Buddenhagen 2006), or, in other words, “search and destroy” activities. In that case, the program arguably skipped delimitation and went straight to control. Additionally, a larger, one-time survey may still be logistically and financially preferable to a dynamic situation in which the total resources that will be required and the duration of the activity are unknown. The DTDS approach was built to focus on a one-shot determination of spatial extent, to facilitate evidence-based planning decisions that lead to management success (Mangel et al. 1984; Panetta and Lawes 2005).

The cost of eradication programs is largely a function of their total area (Woldendorp and Bomford 2004; Harris and Timmins 2009), because often that entire area will be searched (perhaps repeatedly). While boundary accuracy is desirable to limit costs, overestimation is still better than underestimation. Unfortunately, circular areas increase by the square of the radius, so doubling the radius gives four times the area. However, managers can take steps to reduce overestimation if the defined area challenges program capacity. Trying to reduce uncertainty by scouting or increasing the number of positives is one possible response, when resources are available (and assuming variance does not increase). Another option could be choosing a lesser percentile estimate (e.g., 95^th^). This would likely increase the risk of pest egress, but 99^th^ percentile estimates in the examples above were often well beyond the true extent. More generally, any boundary estimate could be viewed as the starting point for a possible quarantine area (containment), subject to revision (reduction) based on further evidence. In other words, bounded areas may not need to be exhaustively searched, depending on management options. More complex methods exist to estimate ranges with uncertainty as polygons (Keith et al. 2019).

## Statements and Declarations

### Data accessibility statement

The novel simulation models created for this research are available upon request from the corresponding author.

## Supplemental materials

### Total transect length calculation

The approach here is similar to an estimate of the required total transect length (*L*_B_) based on the desired coefficient of variation, σ (Buckland et al. 2015):

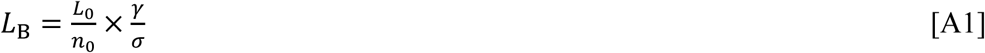

where *L*_0_ is the transect length inspected in pilot tests, *n*_0_ is the number of organisms found, and γ is a shape parameter of the detection function, with a recommended value of 3. The value of *L*_0_/*n*_0_ estimates the reciprocal of baseline *m* (or *m*_0_). If *m* is 0.001 per m, and σ = 0.05, the result is *L*_B_ = 60,000 m.

---

**Figure S1.**
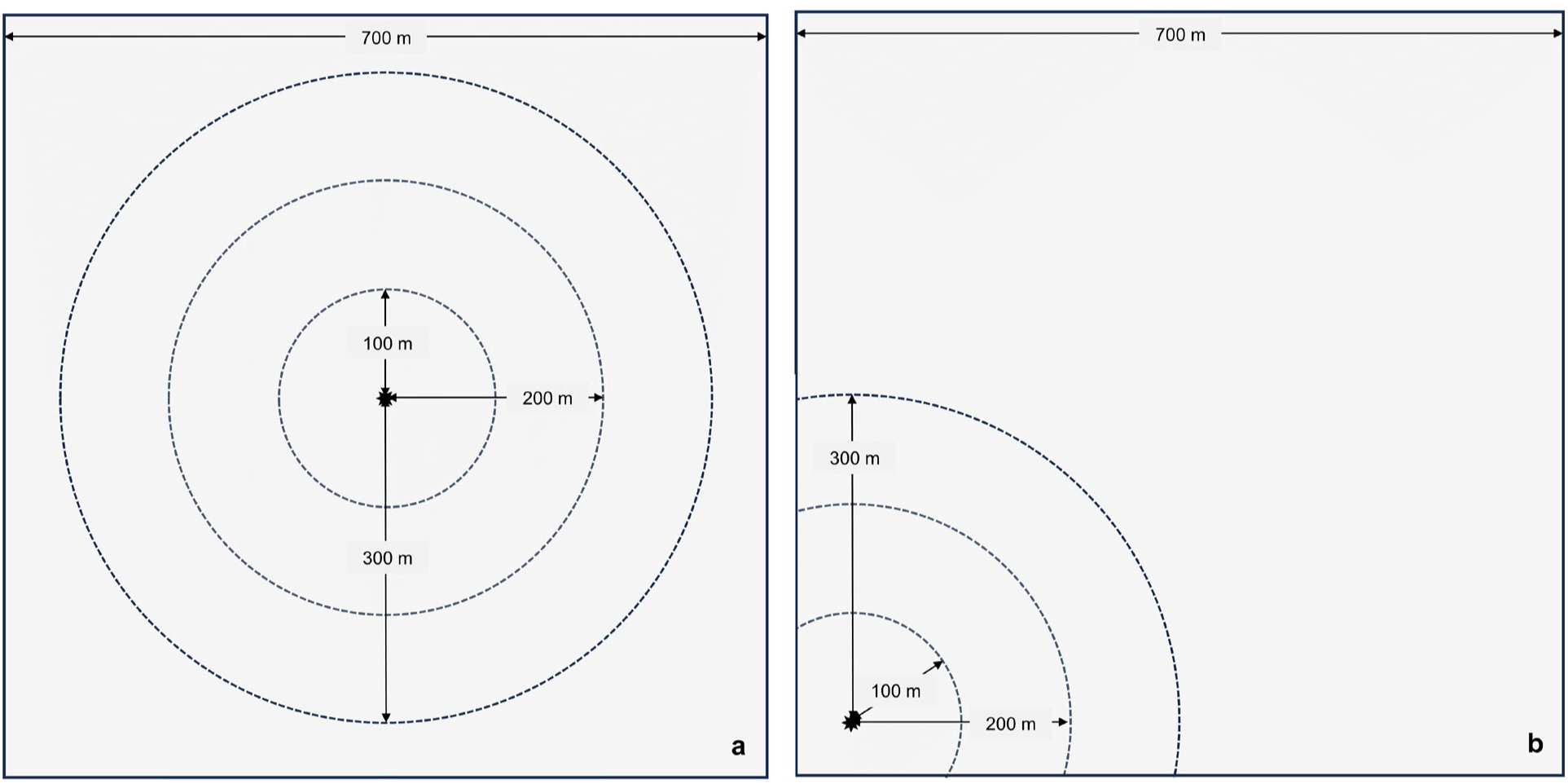
The infestation epicenters and areas of decreasing infestations rates (dashed circles) of Tomato Brown Rugose Fruit Virus (TBRFV) in two case studies: a) centered within the field, or b) near one corner of the field.

**Figure S2.**
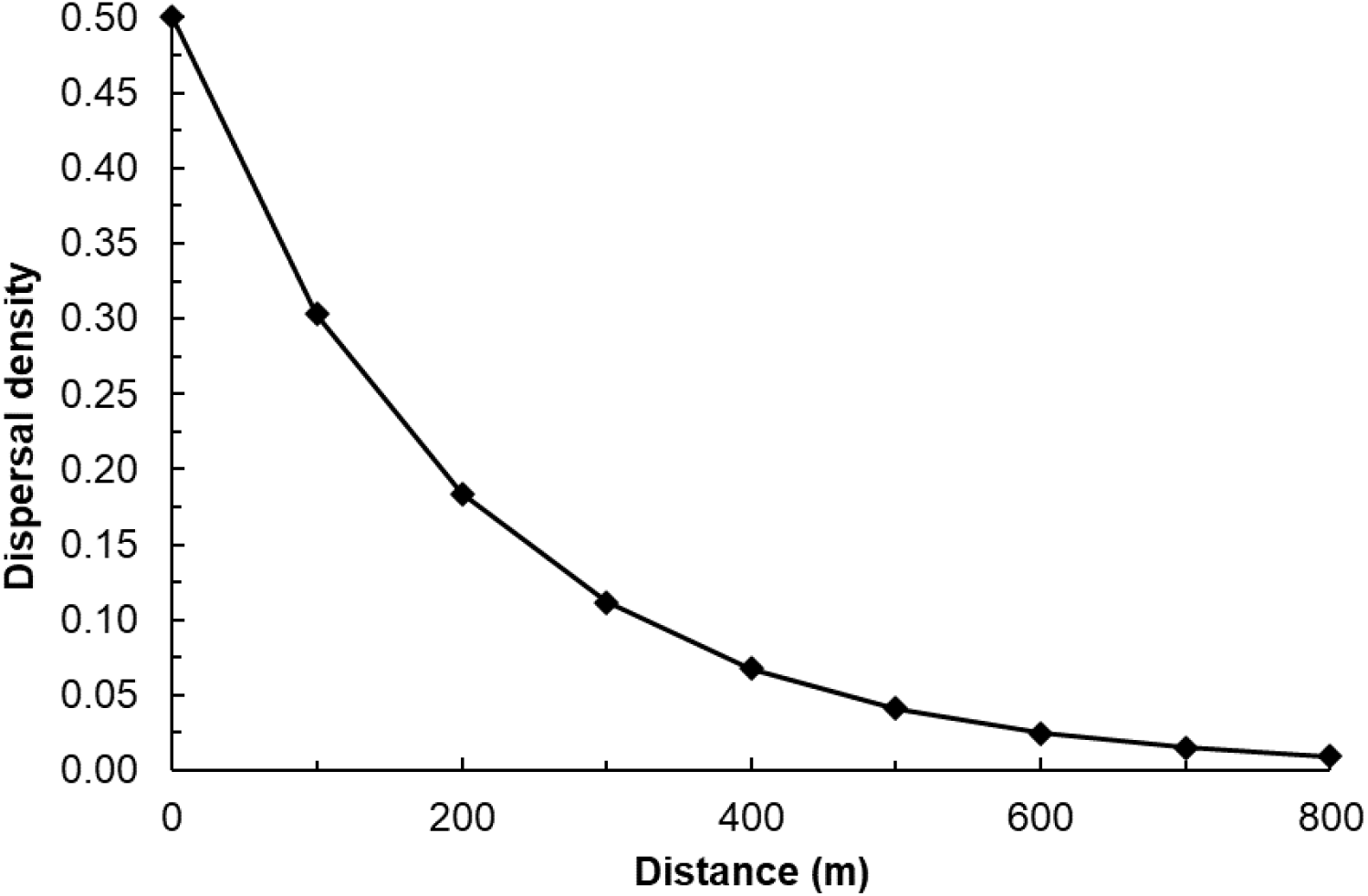
The hypothetical dispersal density for *Phyllosticta citrocarpa* in the Scenario 1 case study.

**Figure S3.**
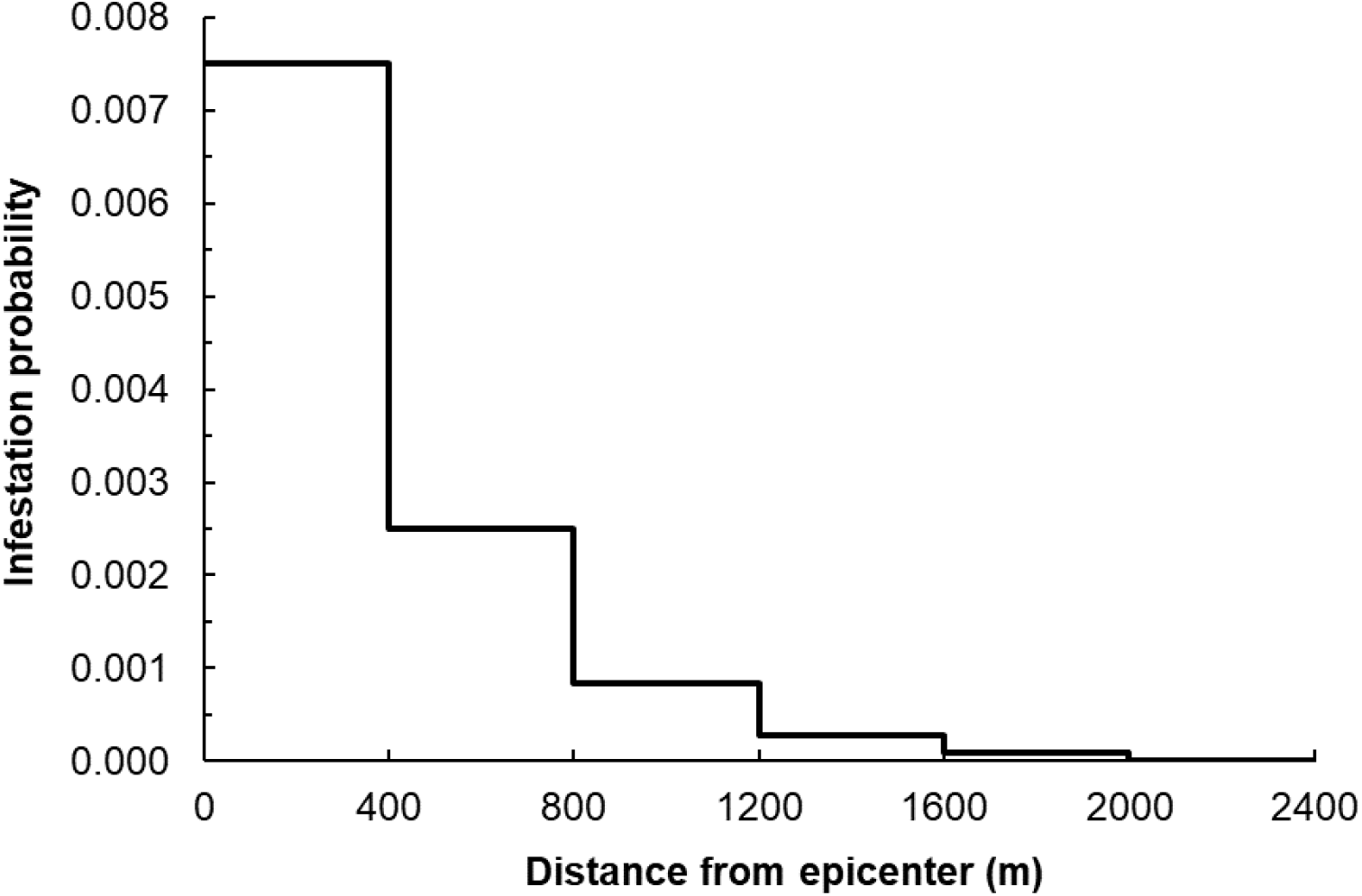
The probability of infestation as a function of distance for *Phyllosticta citrocarpa* in the Scenario 2 case study.

**Figure S4.**
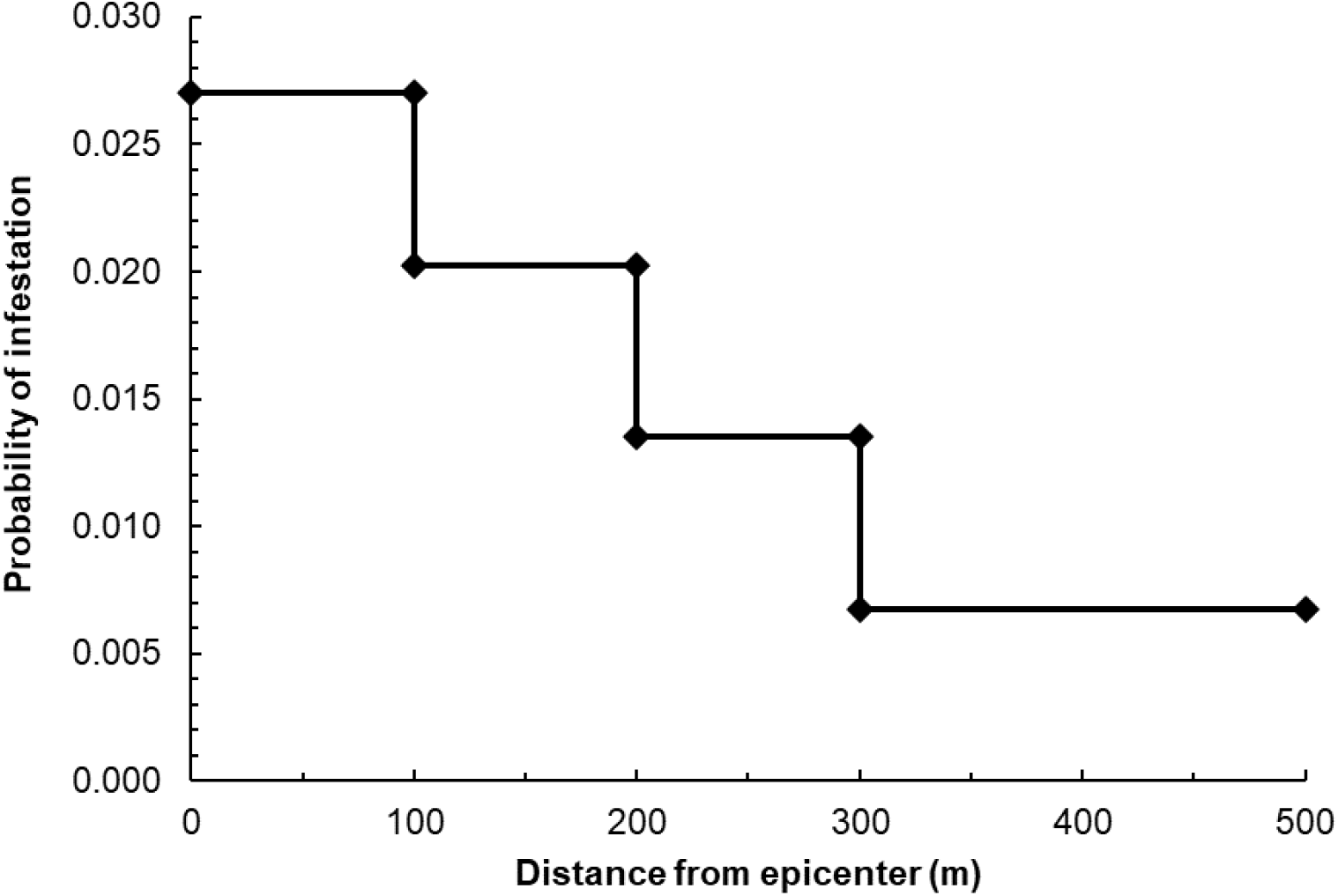
The probability of infestation as a function of distance in the tomato brown rugose fruit virus case study.

**Figure S5.**
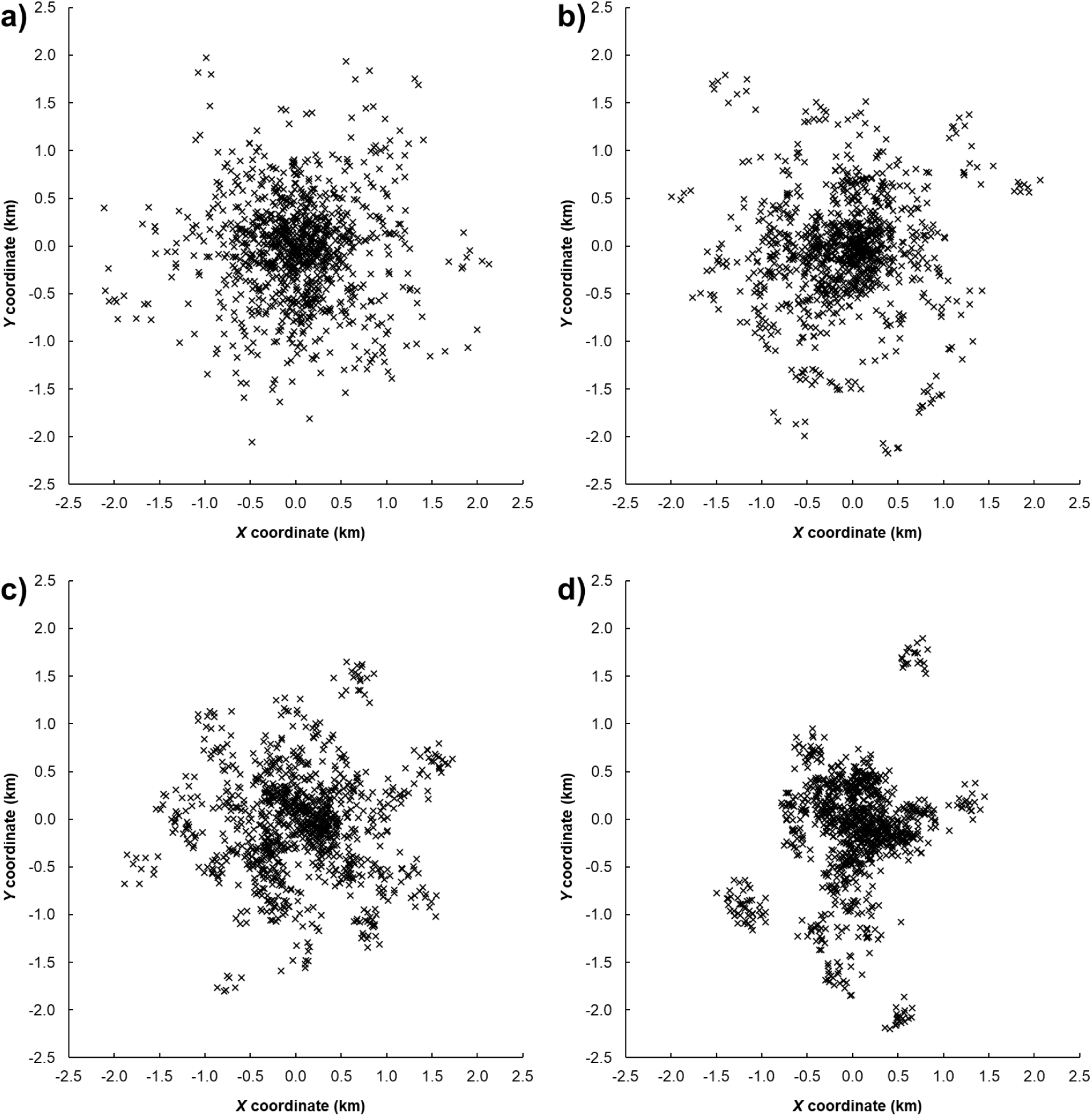
Random positive locations for a less skewed distribution created using a Thomas point process with a values of μ equal to a) 1, b) 6, c) 12, and d) 24 [increasingly clustered].

**Figure S6.**
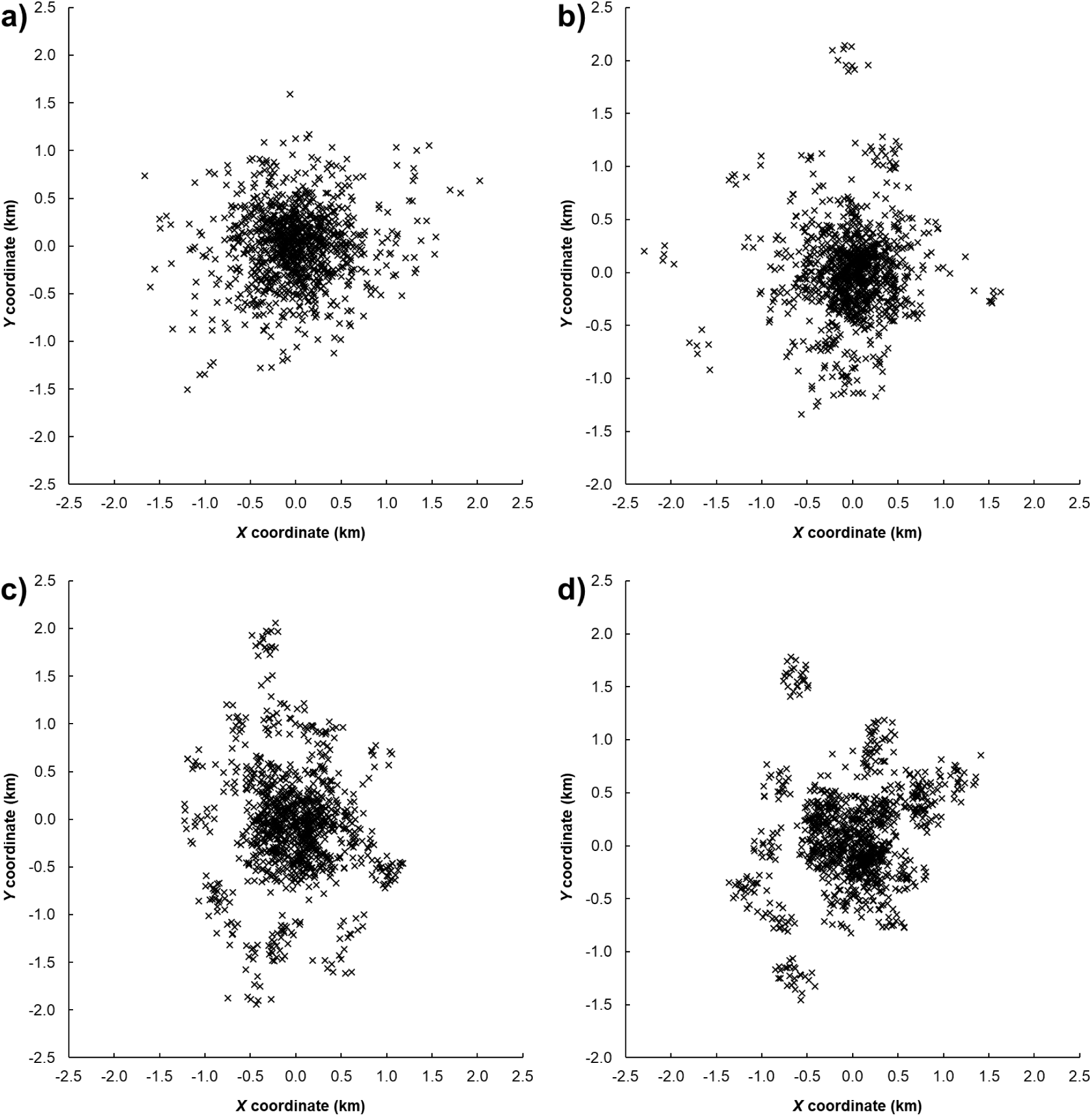
Random positive locations for a more skewed distribution created using a Thomas point process with a values of μ equal to a) 1, b) 6, c) 12, and d) 24 [increasingly clustered].

## REFERENCES

Ahmed DA, Hudgins EJ, Cuthbert RN, Kourantidou M, Diagne C, Haubrock PJ, Leung B, Petrovskii S, Courchamp F. 2022. Managing biological invasions: the cost of inaction. Biological Invasions 24:1927–1946.

Alvarez S, Solís D. 2018. Rapid response lowers eradication costs of invasive species. Choices 33:1–9.

Amburgey SM, Miller DAW, Brand A, Dietrich A, Campbell Grant EH. 2019. Knowing your limits: Estimating range boundaries and co-occurrence zones for two competing plethodontid salamanders. Ecosphere 10:e02727.

Anderson DR, Laake JL, Crain BR, Burnham KP. 1979. Guidelines for line transect sampling of biological populations. The Journal of Wildlife Management:70–78.

Baddeley A, Rubak E, Turner R. 2015. Spatial Point Patterns: Methodology and Applications with R. London: Chapman and Hall/CRC Press.

Barclay HJ. 2021. Mathematical models for using sterile insects. In: Dyck VA, Hendrichs J, Robinson AS, editors. Sterile Insect Technique, Second ed. Boca Raton, FL: CRC Press. p 201–244.

Bate LJ, Torgersen TR, Wisdom MJ, Garton EO, Clabough SC. 2008. Log sampling methods and software for stand and landscape analyses (PNW-GTR-746). In. Portland, Oregon: United States Department of Agriculture, Forest Service, Pacific Northwest Research Station. p 93.

Becker FS, Slingsby JA, Measey J, Tolley KA, Altwegg R. 2022. Finding rare species and estimating the probability that all occupied sites have been found. Ecological Applications 32:e2502.

Bognar M. 2022. Gamma Distribution. In. Ames, Iowa: Department of Statistics and Actuarial Science, University of Iowa.

Bried JT, Pellet J. 2012. Optimal design of butterfly occupancy surveys and testing if occupancy converts to abundance for sparse populations. Journal of Insect Conservation 16:489–499.

Buckland ST, Borchers DL, Johnston A, Henrys PA, Marques TA. 2007. Line transect methods for plant surveys. Biometrics 63:989–998.

Buckland ST, Rexstad EA, Marques TA, Oedekoven CS. 2015. Distance sampling: methods and applications. New York, NY: Springer.

Buddenhagen CE. 2006. The successful eradication of two blackberry species *Rubus megalococcus* and *R. adenotrichos* (Rosaceae) from Santa Cruz Island, Galapagos, Ecuador. Pacific Conservation Biology 12:272–278.

Bullock JM, González LM, Tamme R, Götzenberger L, White SM, Pärtel M, Hooftman DAP. 2017. A synthesis of empirical plant dispersal kernels. Journal of Ecology 105:6–19.

Carpenter SR, Booth EG, Kucharik CJ. 2018. Extreme precipitation and phosphorus loads from two agricultural watersheds. Limnology and Oceanography 63:1221–1233.

Caton BP, Fang H, Manoukis NC, Pallipparambil GR. 2021. Quantifying insect dispersal distances from trapping detections data to predict delimiting survey radii. Journal of Applied Entomology 146:203–216.

CDFA. 2020. Glassy-Winged Sharpshooter Statewide Detection & Delimitation Protocols. In. Sacramento, CA: California Department of Food and Agriculture (CDFA). p 14.

Cochran WG. 1977. Sampling techniques, 3rd ed. New York: Wiley.

Davison AC, Huser R. 2015. Statistics of extremes. Annual Review of Statistics and its Application 2:203–235.

Dodds KJ, Orwig DA. 2011. An invasive urban forest pest invades natural environments—Asian longhorned beetle in northeastern US hardwood forests. Canadian Journal of Forest Research 41:1729–1742.

EFSA, Lázaro E, Parnell S, Civera AV, Schans J, Schenk M, Abrahantes JC, Zancanaro G, Vos S. 2020a. General guidelines for statistically sound and risk-based surveys of plant pests. In. Parma, Italy: European Food Safety Authority (EFSA). p 65.

EFSA, Lázaro E, Parnell S, Civera AV, Schans J, Schenk M, Schrader G, Abrahantes JC, Zancanaro G, Vos S. 2020b. Guidelines for statistically sound and risk-based surveys of Phyllosticta citricarpa. In. Parma, Italy: European Food Safety Authority (EFSA). p 75.

Elzinga CL, Salzer DW. 1998. Measuring & Monitoring Plant Populations. Denver, Colorado: U.S. Department of the Interior, Bureau of Land Management.

Favaro R, Wichmann L, Ravn HP, Faccoli M. 2015. Spatial spread and infestation risk assessment in the Asian longhorned beetle, *Anoplophora glabripennis*. Entomologia Experimentalis et Applicata 155:95–101.

Flesch AD, Murray IW, Gicklhorn JM, Powell BF. 2019. Application of distance sampling for assessing abundance and habitat relationships of a rare Sonoran Desert cactus. Plant Ecology 220:1029–1042.

Forest Service. 2022. Pest Alert: Beech Leaf Disease (R9–PR–001–21). In. Milwaukee, WI: Eastern Region, State and Private Forestry, Forest Service, United States Department of Agriculture. p 2.

Fosgate GT. 2009. Practical sample size calculations for surveillance and diagnostic investigations. Journal of Veterinary Diagnostic Investigation 21:3–14.

García C, Borda-de-Água L. 2017. Extended dispersal kernels in a changing world: insights from statistics of extremes. Journal of Ecology 105:63–74.

Green RH, Young RC. 1993. Sampling to detect rare species. Ecological Applications 3:351–356.

Guillera-Arroita G, Morgan BJT, Ridout MS, Linkie M. 2011. Species occupancy modeling for detection data collected along a transect. Journal Of Agricultural, Biological, and Environmental Statistics 16:301–317.

Harris S, Timmins SM. 2009. Estimating the benefit of early control of all newly naturalised plants. Science for Conservation 292. In: Science for Conservation 292. Wellington, New Zealand: Department of Conservation.

Hauser CE, McCarthy MA. 2009. Streamlining ‘search and destroy’: cost-effective surveillance for invasive species management. Ecology Letters 12:683–692.

Henderson DC. 2010. Occupancy survey guidelines for prairie plant species at risk. In. Saskatoon, Saskatchewan: Prairie and Northern Region, Canadian Wildlife Service, Environment Canada. p 45.

Herrick JE, Zee JWV, Havstad KM, Burkett LM, Whitford WG. 2005. Monitoring Manual for Grassland, Shrubland and Savanna Ecosystems. Volume I: Quick Start. In. Las Cruces, New Mexico: USDA - ARS Jornada Experimental Range. p 42.

Hester SM, Hauser CE, Kean JM. 2017. Tools for Designing and Evaluating Post-Border Surveillance Systems. In: Robinson AP, Walshe T, Burgman MA, Nunn M, editors. Invasive species: Risk assessment and management. Cambridge, United Kingdom: Cambridge University Press. p 17–52.

Hilborn R, Mangel M. 1997. The Ecological Detective: Confronting Models with Data. Princeton, NJ: Princeton University Press.

Howell CJ. 2012. Progress toward environmental weed eradication in New Zealand. Invasive Plant Science and Management 5:249–258.

Hull-Sanders H, Pepper E, Davis K, Trotter III RT. 2017. Description of an establishment event by the invasive Asian longhorned beetle (*Anoplophora glabripennis*) in a suburban landscape in the northeastern United States. PLoS One 17:e0181655.

IPPC. 2016. International Standards for Phytosanitary Measures, Publication No. 5. Glossary of Phytosanitary Terms. . In. Rome, Italy: International Plant Protection Convention (IPPC), Food and Agriculture Organization of the United Nations. p 38.

IPPC. 2021. Surveillance Guide: A guide to understand the principal requirements of surveillance programmes for national plant protection organizations (Second Edition). In, Second ed. Rome, Italy. http://www.fao.org/3/cb7139en/cb7139en.pdf: International Plant Protection Convention (IPPC), Food and Agriculture Organization of the United Nations. p 76.

Jacobs LEO, Richardson DM, Wilson JRU. 2014. *Melaleuca parvistaminea* Byrnes (Myrtaceae) in South Africa: invasion risk and feasibility of eradication. South African Journal of Botany 94:24–32.

James K, Bradshaw K. 2020. Detecting plant species in the field with deep learning and drone technology. Methods in Ecology and Evolution 11:1509–1519.

Jang EB, Enkerlin W, Miller C, Reyes-Flores J. 2014. Trapping Related to Phytosanitary Status and Trade. In: Shelly T, Epsky N, Jang EB, Reyes-Flores J, Vargas R, editors. Trapping and the Detection, Control, and Regulation of Tephritid Fruit Flies. Dordrecht: Springer. p 589–608.

Jiménez MA, Jaksic FM, Armesto JJ, Gaxiola A, Meserve PL, Kelt DA, Gutiérrez. JR. 2011. Extreme climatic events change the dynamics and invasibility of semi-arid annual plant communities. Ecology Letters 4:12227–11235.

Jones O, Maillardet R, Robinson A. 2009. Introduction to Scientific Programming and Simulation Using R. New York: Chapman and Hall/CRC.

Kalischuk M, Paret ML, Freeman JH, Raj D, Da Silva S, Eubanks S, Wiggins DJ, Lollar M, Marois JJ, Mellinger HC, Das J. 2019. An improved crop scouting technique incorporating unmanned aerial vehicle–assisted multispectral crop imaging into conventional scouting practice for gummy stem blight in watermelon. Plant Disease 103:1642–1650.

Katz RW, Brush GS, Parlange MB. 2005. Statistics of extremes: modeling ecological disturbances. Ecology 86:1124–1134.

Kean JM, Burnip GM, Pathan A. 2015. Detection survey design for decision making during biosecurity incursions. In: Jarrad F, Low-Choy S, Mengersen K, editors. Biosecurity Surveillance: Quantitative Approaches. Boston, MA: CABI. p 238–252.

Keith DA. 2000. Sampling designs, field techniques and analytical methods for systematic plant population surveys. Ecological Management & Restoration 1:125–139.

Keith JM, Spring D, Kompas T. 2019. Delimiting a species’ geographic range using posterior sampling and computational geometry. Scientific Reports 9:1–15.

Koh JMJ, Cunniffe NJ, Parnell S. 2025. Assessing delimiting strategies to identify the infested zones of quarantine plant pests and diseases. Scientific Reports 15:5610.

Lawes R, Panetta FD. 2004. Detecting alien plant species early in the invasion process: a sampling strategy for the detection of Chromolaena odorata (L.) RM King and H. Rob.(Siam weed). In: 14th Australian Weeds Conference: Weed management: balancing people, planet, profit. Wagga Wagga, New South Wales, Australia: Weed Society of New South Wales. p 484–487.

Lázaro E, Sesé M, López-Quílez A, Conesa D, Dalmau V, Ferrer-Matoses A, Vicent A. 2020. Tracking the outbreak. An optimized delimiting survey strategy for Xylella fastidiosa. bioRxiv:58 pp.

Leung B, Cacho OJ, Spring D. 2010. Searching for non-indigenous species: rapidly delimiting the invasion boundary. Diversity and Distributions 16:451–460.

MacKenzie DI, Royle JA, Brown JA, Nichols JD. 2013. Occupancy Estimation and Modeling for Rare and Elusive Populations. In: Thompson W, editor. Sampling Rare or Elusive Species: Concepts, Designs, and Techniques for Estimating Population Parameters. Washington, D.C.: Island Press. p 149–172.

Mangel M, Plant RE, Carey JR. 1984. Rapid delimiting of pest infestations: a case study of the Mediterranean fruit fly. Journal of Applied Ecology 21:563–579.

McGarvey R, Burch P, Matthews JM. 2016. Precision of systematic and random sampling in clustered populations: habitat patches and aggregating organisms. Ecological Applications 26:233–248.

Meats A. 1998a. Cartesian methods of locating spot infestations of the papaya fruit fly ‘*Bactrocera papayae*’ Drew and Hancock within the trapping grid at Mareeba, Queensland, Australia. General and Applied Entomology: The Journal of the Entomological Society of New South Wales 28:57–60.

Meats A. 1998b. The power of trapping grids for detecting and estimating the size of invading propagules of the Queensland fruit fly and risks of subsequent infestation. General and Applied Entomology 28:47–55.

Meats A, Clift AD, Robson MK. 2003. Incipient founder populations of Mediterranean and Queensland fruit flies in Australia: the relation of trap catch to infestation radius and models for quarantine radius. Australian Journal of Experimental Agriculture 43:397–406.

NASS. 2019. 2017 Census of Agriculture, California State and County data. In: Geographic Area Series. Washington, D.C.: United States Department of Agriculture, National Agricultural Statistics Service (NASS). p 720.

Nathan R, Klein EK, Robledo-Arnuncio JJ, Revilla E. 2012. Dispersal Kernels: Review. In: Clobert J, Baguette M, Benton TG, Bullock JM, editors. Dispersal Ecology and Evolution Oxford University Press. p 187–210.

Nomani SZ, Oli MK, Carthy RR. 2012. Line transects by design: The influence of study design, spatial distribution and density of objects on estimates of abundance. Open Ecology Journal 5:25–44.

Panetta FD, Lawes R. 2005. Evaluation of weed eradication programs: the delimitation of extent. Diversity and Distributions 11:435–442.

Parkes JP, Panetta FD. 2009. Eradication of invasive species: progress and emerging issues in the 21st century. In: Clout MN, Williams PA, editors. Invasive species management. A handbook of principles and techniques. Oxford, U.K.: Oxford University Press. p 47–60.

PDA. 2022. Pennsylvania Spotted Lanternfly Program. In. Harrisburg, PA: Pennsylvania Department of Agriculture (PDA).

Perryman SAM, Clark SJ, West JS. 2014. Splash dispersal of *Phyllosticta citricarpa* conidia from infected citrus fruit. Scientific Reports 4:6568.

Philippi T. 2005. Adaptive cluster sampling for estimation of abundances within local populations of low-abundance plants. Ecology 86:1091–1100.

Potts JM, Cox MJ, Barkley P, Christian R, Telford G, Burgman MA. 2013. Model-based search strategies for plant diseases: a case study using citrus canker (*Xanthomonas citri*). Diversity and Distributions 19:590–602.

PPQ. 2014. Asian Longhorned Beetle Response Guidelines. In. Raleigh, NC: U.S. Department of Agriculture (USDA), Animal and Plant Health Inspection Service (APHIS), Plant Protection and Quarantine (PPQ). p 17.

PPQ. 2018. New Pest Response Guidelines,Cucumber green mottle mosaic virus CGMMV. In. Raleigh, NC: U.S. Department of Agriculture (USDA), Animal and Plant Health Inspection Service (APHIS), Plant Protection and Quarantine (PPQ). p 39.

PPQ. 2020a. ALB [Asian longhorned beetle] Survey Protocol. In. Washington, D.C.: U.S. Department of Agriculture (USDA), Animal and Plant Health Inspection Service (APHIS), Plant Protection and Quarantine (PPQ). p 17.

PPQ. 2020b. New Pest Response Guidelines, Tobamovirus: Tomato brown rugose fruit virus. In. Raleigh, NC: U.S. Department of Agriculture (USDA), Animal and Plant Health Inspection Service (APHIS), Plant Protection and Quarantine (PPQ). p 45.

PPQ. 2022a. 2022 User Manual: Nationwide Visual. In. Riverdale, MD: Plant Protection and Quarantine (PPQ), Animal and Plant Health Inspection Service, United States Department of Agriculture.

PPQ. 2022b. New Pest Response Guidelines: Cydalima perspectalis, Box tree moth. In. Washington, D.C.: U.S. Department of Agriculture, Animal and Plant Health Inspection Service, Plant Protection and Quarantine (PPQ). p 54.

Reiss R-D, Thomas M. 2007. Statistical Analysis of Extreme Values: with Applications to Insurance, Finanace, Hydrology and Other Fields (3rd Edition). Boston: Birkhäuser.

Ringvall A. 2007. New methods for sampling sparse populations. In: McRoberts RE, Reams GA, Van Deusen PC, McWilliams WH, editors. Seventh Annual Forest Inventory and Analysis Symposium. Portland, ME: US Department of Agriculture, Forest Service. p 215–220.

Roberts J, Low-Choy S, Jarrad F, Mengersen K. 2015. Common Statistical Distributions Used in Statistical Modelling and Analysis for Biosecurity Surveillance. In: Jarrad F, Low-Choy S, Mengersen K, editors. Biosecurity Surveillance: Quantitative Approaches. Boston, MA: CABI. p 348–361.

Schroeder M. 2012. Strategies for detection and delimitation surveys of the pine wood nematode in Sweden. In. Jönköping, Sweden: Swedish Board of Agriculture, for the Department of Ecology at the Swedish University of Agricultural Sciences. p 36.

Scott DW. 2009. Sturges’ rule. Wiley Interdisciplinary Reviews: Computational Statistics 1:303–306.

Scott JK, Batchelor KL. 2014. Management of *Chrysanthemoides monilifera* subsp. *rotundata* in Western Australia. Invasive Plant Science and Management 7:190–196.

Simberloff D. 2003. Eradication-preventing invasions at the outset. Weed Science 51:247–253.

Skarpaas O, Shea K. 2007. Dispersal patterns, dispersal mechanisms, and invasion wave speeds for invasive thistles. The American Naturalist 170:421–430.

Smith T, Whilby L, Derksen A. 2010. Florida CAPS/DPI Giant African Snail, Achatina spp. (Pulmonata: Achatinidae) Survey Report No. 2010-02-GAS-01. In. Gainesville, FL: Division of Plant Industry, Florida Department of Agriculture and Consumer Services. p 14.

Sun H, Douma JC, Schenk MF, van der Werf W. 2025. Comparing inward and outward strategies for delimiting non-native plant pest outbreaks. Journal of Pest Science 10:1–6.

Thompson W, editor. 2013. Sampling rare or elusive species: concepts, designs, and techniques for estimating population parameters. Washington, D.C.: Island Press.

Torres-Quezada E, Gandini-Taveras RJ. 2023. Plant Density Recommendations and Plant Nutrient Status for High Tunnel Tomatoes in Virginia. Horticulturae 9:1063.

Turgeon JJ, Orr M, Grant C, Wu Y, Gasman B. 2015. Decade-Old Satellite Infestation of *Anoplophora glabripennis* Motschulsky (Coleoptera: Cerambycidae) Found in Ontario, Canada Outside Regulated Area of Founder Population. The Coleopterists Bulletin 69:674–678.

Turgeon JJ, Pedlar J, De Groot P, Smith MT, Jones C, Orr M, Gasman B. 2010. Density and location of simulated signs of injury affect efficacy of ground surveys for Asian longhorned beetle. The Canadian Entomologist 142:80–96.

USFS Pacific Northwest Region. 2012. Stream Inventory Handbook Level I & II. In. Portland, Oregon: Department of Agriculture, United States Forest Service (USFS), Pacific Northwest Region. p 127.

van Havre Z, Whittle P. 2015. Designing Surveillance for Emergency Response. In: Jarrad F, Low-Choy S, Mengersen K, editors. Biosecurity Surveillance: Quantitative Approaches. Boston, MA: CABI. p 123–133.

Vose D. 2000. Risk Analysis: A Quantitative Guide, 2nd ed. New York: John Wiley & Sons, Ltd.

Weldon CW, Schutze MK, Karsten M. 2014. Trapping to monitor tephritid movement: results, best practice, and assessment of alternatives. In: Shelly T, Epsky N, Jang EB, Reyes-Flores J, Vargas R, editors. Trapping and the Detection, Control, and Regulation of Tephritid Fruit Flies. Dordrecht: Springer. p 175–217.

Welsh MJ, Turner JA, Epanchin-Niell RS, Monge JJ, Soliman T, Robinson AP, Kean JM, Phillips C, Stringer LD, Vereijssen J, Liebhold AM. 2021. Approaches for estimating benefits and costs of interventions in plant biosecurity across invasion phases. Ecological Applications 31:e02319.

Woldendorp G, Bomford M. 2004. Weed eradication: strategies, timeframes and costs. In. Canberra, Australia: Bureau of Resource Sciences, Department of Agriculture, Fisheries and Forestry. p 30.

